# Elucidating Mechanisms of Tolerance to *Salmonella* Typhimurium Across Long-Term Infections Using the Collaborative Cross

**DOI:** 10.1101/2022.04.11.487981

**Authors:** Kristin Scoggin, Jyotsana Gupta, Rachel Lynch, Aravindh Nagarajan, Manuchehr Aminian, Amy Peterson, L. Garry Adams, Michael Kirby, David W. Threadgill, Helene L. Andrews-Polymenis

## Abstract

Understanding the molecular mechanisms underlying resistance and tolerance to pathogen infection may present the opportunity to develop novel interventions. Resistance is the absence of clinical disease with low pathogen burden, while tolerance is minimal clinical disease in the face of high pathogen burden. *Salmonella* is a worldwide health concern. We studied 18 strains of Collaborative Cross mice that survive acute *Salmonella* Typhimurium (STm) infections. We infected these strains orally and monitored them for three weeks post-infection. Five strains cleared STm by the end of the experiment (resistant), while 6 strains maintained a bacterial load and survived to the end of the experiment (tolerant). The remaining 7 strains survived longer than 7 days but succumbed to infection before the end of the study period and were called “delayed susceptible” to differentiate them from strains that do not survive to day 7 (susceptible). Tolerant strains were colonized in Peyer’s patches, mesenteric lymph node, spleen and liver, while resistant strains had significantly reduced bacterial colonization. Tolerant strains had lower pre-infection core body temperatures than both delayed susceptible and resistant strains and had disrupted circadian patterns of body temperature post-infection sooner than resistant strains. Tolerant strains had higher circulating total white blood cells than resistant strains, driven by increased numbers of neutrophils. Tolerant strains had more severe tissue damage and higher circulating levels of MCP-1 and IFN-γ, but lower levels of ENA-78 than resistant strains. QTL analysis revealed 1 significant association and 6 suggestive associations. RNA-seq analysis identified 22 genes that are differentially regulated in tolerant versus resistant animals that overlapped with the QTLs we identified and allowed us to identify the top 5 canonical pathways. Fibrinogen genes (*Fga*, *Fgb*, and *Fgg*) were found across the QTL, RNA, and top canonical pathways making them the best candidate genes for differentiating tolerance and resistance.

**Author Summary:** An infected host can respond in multiple ways to bacterial infection including resistance and tolerance. Resistance is a decrease in pathogen load, while in tolerance mild clinical signs are present despite high pathogen load. We infected a collection of 18 strains of genetically diverse mice with *Salmonella* Typhimurium for up to three weeks. Five strains were resistant, 6 strains were tolerant, and the remaining 7 strains survived an intermediate amount of time (“delayed susceptible”). Tolerant strains maintained bacterial load across several organs, while resistant strains reduced bacterial load. Tolerant strains had the lowest pre-infection core body temperatures and the most rapid disruption in circadian patterns of body temperature post-infection. Tolerant strains had higher circulating neutrophils, higher circulating levels of MCP-1 and IFN-γ, but lower levels of ENA-78 than resistant strains, in addition to more severe tissue damage than resistant strains. QTL analysis revealed multiple associated regions, and gene expression analysis identified 22 genes that are differentially regulated in tolerant versus resistant animals in these regions. Fibrinogen genes (*Fga*, *Fgb*, and *Fgg*) were found across the QTL, RNA, and top canonical pathways making them the best candidate genes for differentiating tolerance and resistance.

## Introduction

*Salmonella enterica* are Gram negative bacteria that cause a range of diseases including gastroenteritis, sepsis, and typhoid fever in various mammalian and avian hosts [1, 2]. 93.8 million cases Non-typhoidal *Salmonella* (NTS) occur world-wide every year in humans, resulting in 155,000 deaths [3]. The severity of an NTS infection depends on several factors including host age, health, and genetics, as well as *Salmonella* serotype [4–6]. *Salmonella enterica* serotype Typhimurium (STm) causes gastroenteritis in humans and can cause serious bacteremia. In mice, this serotype causes a fatal systemic infection in susceptible mouse strains that models invasive infections in humans [1, 7].

Various strains of inbred mice respond to STm infection differently. C57BL/6J and BALB/cJ have mutations in solute carrier family 11 member 1 (*Slc11a1*) and are highly susceptible to fatal STm bacteremia [8–10]. Other strains including 129Sv, CBA/J, and A/J, are wild type at the *Slc11a1* locus and do not develop severe systemic salmonellosis [8–11]. Other genes, including neutrophil cytosolic factor 2 (*Ncf2*), toll-like receptor 4 (*Tlr4*), interferon gamma (*Ifng*), and histocompatibility complex (H2) haplotypes have also been linked to survival after STm infections in mice [1,12–15].

Since previous STm infection studies have used primarily traditionally inbred mouse strains, the host genetic diversity influencing STm infection in the mouse has not been thoroughly explored. The Collaborative Cross (CC) mouse population is a panel of recombinant inbred strains that recapitulate the genetic diversity found in human populations [16–19]. The CC founders capture 90% of the genetic diversity found in the mouse genome and represents a wider range of phenotypes, including immune phenotypes, than traditional inbred strains [20–22]. For example, strains with high viral titers after experimental infection with influenza virus or Ebola virus and good health status have been identified in the CC [23–25].

How a host responds to infection can be categorized as resistant or susceptible, based on the host’s ability to clear the infection [1, 26]. A third response called tolerance has been described in plant systems and is now being applied to other systems including *Drosophila* and mice [27–33]. In tolerance, the host has minimal signs of infection in the face of a high pathogen burden [32,34,35] and this quality has been attributed to the ability to limit damage [31]. Genetic variation in both resistance and tolerance has been demonstrated using murine infections with plasmodium [31, 36]. Our recent work with CC mice suggests that while there are multiple genes known to influence acute survival after STm infection, there are additional genes to be discovered that influence disease outcome after STm infection [37]. Studies using the CC mouse population suggest that the disease outcomes after STm infection in mice are much more complex than a binary susceptible and resistant classification [37].

To differentiate tolerance from resistance to *Salmonella* infections, 18 strains of CC mice that survived acute infection were orally infected with STm and monitored for up to three weeks. Our experiments revealed that some CC strains are tolerant to STm infection, and that these strains maintain a significantly higher bacterial burden than resistant strains in Peyer’s Patches (PP), mesenteric lymph nodes (MLN), spleen, and liver. Tolerant strains lost more weight than resistant strains despite surviving infection. Not surprisingly, resistant strains had significantly reduced tissue damage in spleen and liver relative to tolerant strains. Tolerant strains had more circulating total white blood cells, driven by higher circulating neutrophils, and higher serum interferon-gamma (IFN-γ), and monocyte chemoattractant protein-1 (MCP-1) than resistant strains. Resistant strains had significantly more circulating epithelial neutrophil-activating protein 78 (ENA-78) than tolerant strains. Using quantitative trait loci (QTL) analysis, we identified one significant association, *Scq1*, and six suggestive associations. RNA-seq analysis allowed us to narrow these regions to 22 candidate genes and the most highly differentially expressed were the fibrinogen subunits *Fga*, *Fgb*, and *Fgg*. RNA-seq analysis also allowed us to identify the top 5 canonical pathways, which also contained these fibrinogen genes.

## Results

### Three week *Salmonella* infections distinguish tolerant and resistant strains and reveal another phenotype: delayed susceptibility

Based on survival of acute STm infections (7 days), 18 CC mouse strains were chosen to undergo a three-week infection with STm ATCC14028 to differentiate between tolerant and resistant phenotypes. Tolerance was defined as strains that had at least 4/6 mice survive to day 7 during one week infections [37], at least 4/ 6 mice survive to day 21 during three week infections, and had a median colonization of at least 10^3^ CFU/g in liver and 10^4^ CFU/g in spleen (Fig 1A-D). Resistant strains had the same or better survival as tolerant mice but had reduced bacterial burden with a median of <10^3^ CFU/g in liver and <10^4^ CFU/g in spleen (Fig 1A-D).

**Figure 1:**
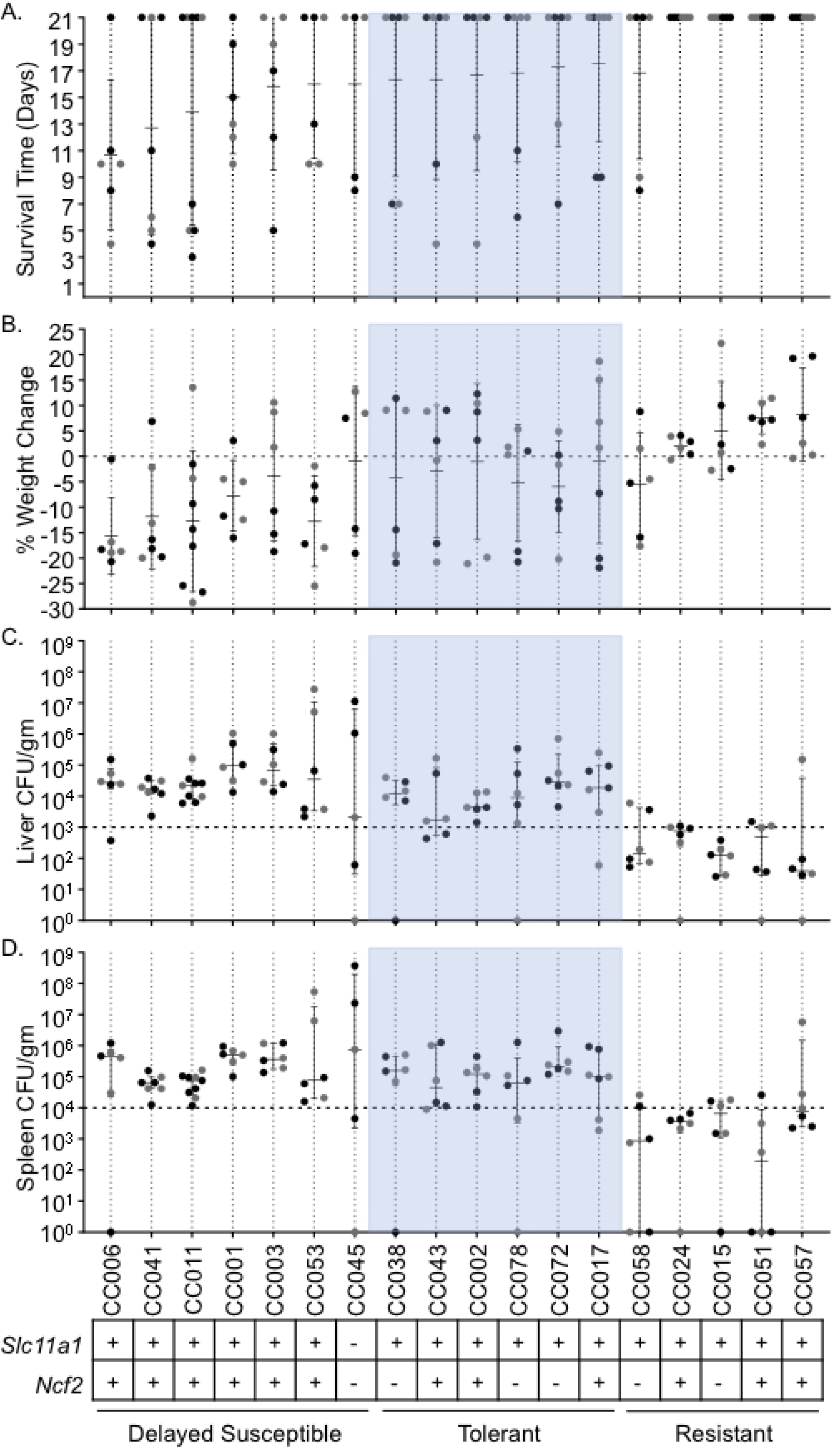
Delayed susceptible, tolerant, and resistant strains have distinct responses to STm infections. A. Survival time and B. percent weight change after infection with STm. Strains are shown in ascending order of survival and weight change if survival was equal between strains. Mean and standard deviation are shown. C. Liver and D. Spleen CFU per gram of organ at time of necropsy. Median and interquartile range shown. Circles represent individual mice (black circles represent males and grey circles represent females). 18 CC strains are represented. *Slc11a1* and *Ncf2* status (+ is wild type, - is mutated) shown at bottom as well as response to infection (tolerant shown in blue).

Of the 18 strains analyzed (see Table S5 for complete strain names), six were categorized as tolerant (CC002, CC017, CC038, CC043, CC072, CC078) and five were categorized as resistant (CC015, CC024, CC051, CC057, CC058). The remaining seven strains did not fit into either the tolerant or resistant category because 3/6 or fewer mice survived the three week study period, despite having at least 4/6 mice survive our earlier acute study [37]. We categorized these strains as “delayed susceptible” since they survive longer than susceptible strains.

Tolerant strains had a mean survival time of 16-18 days (Fig 1A). Four of the five resistant strains had all 6 mice survive to day 21. For the remaining resistant strain, CC058, 4/6 mice survive to day 21 with a mean survival of 16.83 days (Fig 1A). Delayed susceptible strains had a mean survival time of 7-16 days (Fig 1A). In general, both delayed susceptible and tolerant strains lost weight, while resistant strains gained weight. Delayed susceptible strains lost 0.93% - 15.67% of their starting weights while tolerant strains lost 1.02% - 5.97% (Fig 1B). Resistant strains gained 2.05% - 8.17% of their starting weight, with the exception of one strain (CC058 lost 5.51%) (Fig 1B).

We defined tolerance and resistance by a strain’s ability to control the bacterial burden in spleen and liver. Resistant strains ranged in median colonization from 3.9 x10^1^ - 7.53 x10^2^ CFU/g in liver and 1.88 x10^2^ −7.48 x10^3^ CFU/g in spleen (Fig 1C and 1D). Tolerant strains ranged in median colonization from 1.74 x10^3^ −2.78 x10^4^ CFU/g in liver and 4.5 x10^4^ −2.13 x10^5^ CFU/g in spleen (Fig 1C and 1D). Delayed susceptible strains overlapped with the tolerant median colonization ranges for both organs, 2.07 x10^3^ −9.44 x10^4^ CFU/g in liver and 6.5 x10^4^ −7.5 x10^5^ CFU/g in spleen (Fig 1C and 1D). Overlapping ranges of colonization of tolerant and delayed susceptible strains demonstrate that tolerant strains survive a bacterial burden that is fatal for other CC strains. One tolerant strain, CC072, had the highest colonization of all tolerant strains in both spleen and liver, while another, CC043, was more poorly colonized in both organs, illustrating that there is a broad spectrum of bacterial colonization that is tolerated across host genetics.

### Tolerant strains maintain a higher bacterial load in Peyer’s Patches and mesenteric lymph nodes than resistant strains

While our initial focus was on spleen and liver, the ileum, cecum, colon, MLN, and PP were also examined for bacterial load after STm infection. Bacterial burden in ileum, cecum, and colon was not significantly different in tolerant or resistant strains at one or three weeks post-infection (Fig S1, *P* = 0.5600, >0.9999, and >0.9999). Tolerant and resistant strains were colonized similarly in their PP and MLN during acute (one week) infections in previous experiments (Fig 2A and 2B, *P* = >0.9999) [37]. At three week post-infection, tolerant strains remained more highly colonized than resistant strains in both PP (1.28 x10^5^ vs. 1.28 x10^4^ CFU/g, *P* = 0.038) and MLN (6.00 x10^3^ vs. 3.87 x10^2^ CFU/g, *P* = 0.0008) (Fig 2A and 2B). Tolerant strains were stably colonized in PP and MLN between one and three weeks post-infection, suggesting that they were unable to clear STm from these niches (Fig 2A and 2B). Resistant strains however, reduced PP colonization significantly between one and three weeks post-infection (Fig 2A, 1.56 x10^5^ CFU/g and 1.28 x10^4^ CFU/g, *P* = 0.0451), while MLNs remained stably colonized. Thus, during longer-term infections, resistant strains reduce STm colonization of the PP while tolerant strains do not, and resistant strains maintain low levels of MLN colonization.

**Figure 2:**
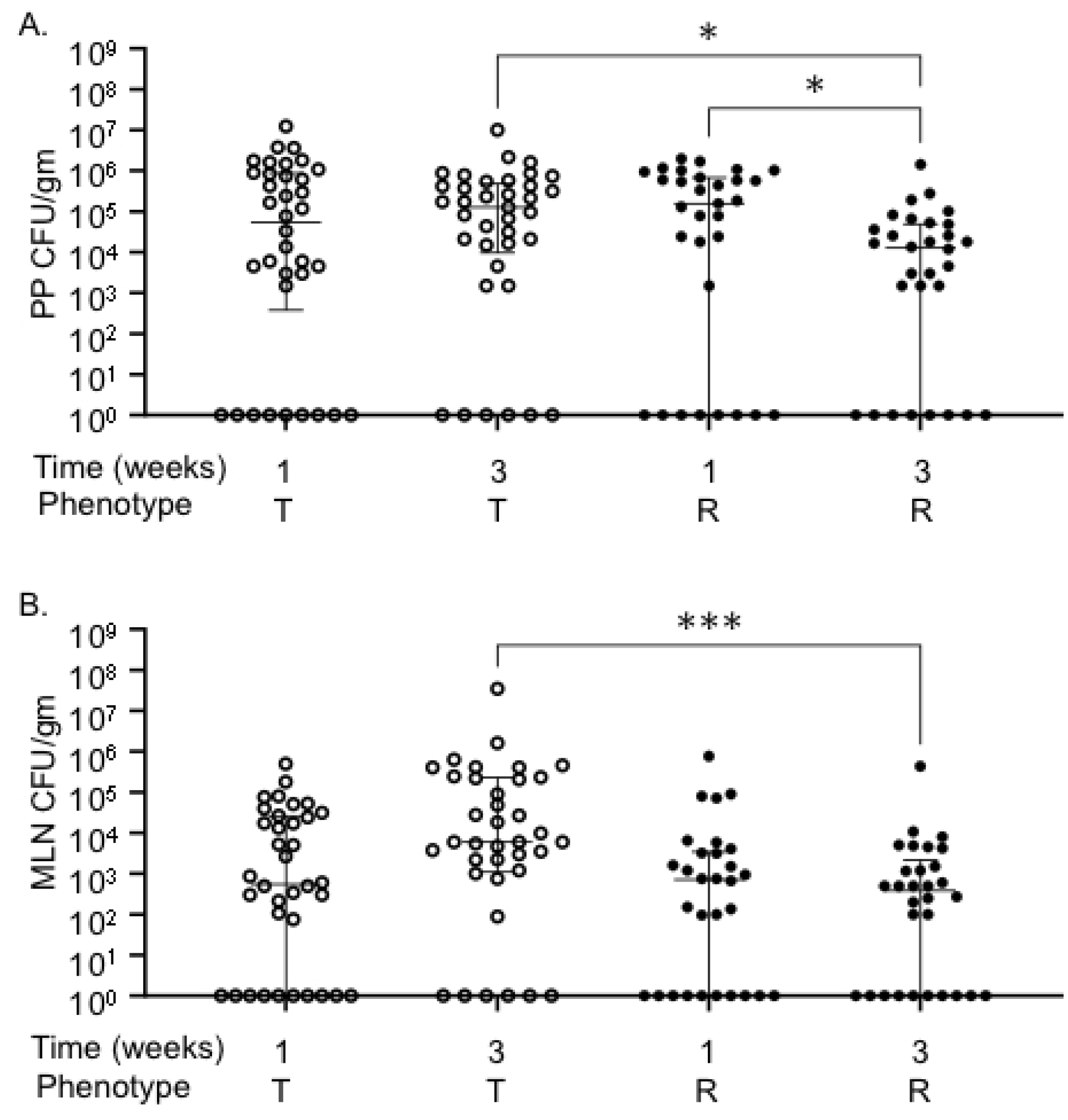
CC strains resistant to STm infection have lower MLN and PP colonization compared to tolerant strains. A. Peyer’s Patches (PP) and B. mesenteric lymph nodes (MLN) CFU per gram of organ at necropsy. Each circle represents an individual mouse and medians and interquartile range are indicated. Tolerant (T, open circles) and resistant (R, filled circles) indicated. Kruskal-Wallis test was performed to find significant differences (* *P* <0.05, ** *P* <0.01, *** *P* <0.001).

### Tolerant strains have lower baseline core body temperature and reduced time to deviation from normal circadian pattern

Mice were implanted with telemetry devices that tracked temperature and activity continuously. Baseline measurements were taken for one week pre-infection and for up to three weeks post-infection. The body temperature minimum corresponds to the minimum core body temperature reached during the “rest period” of the mouse (light hours), the maximum corresponds to the maximum core body temperature reached during the “active period” (dark hours), and the median describes the median core body temperature in a 24-hour period. Tolerant strains had significantly lower baseline body temperatures than both resistant and delayed susceptible strains across all three measurements periods (Fig 3A and S1 Table, rest, active, and 24-hour period).

**Figure 3:**
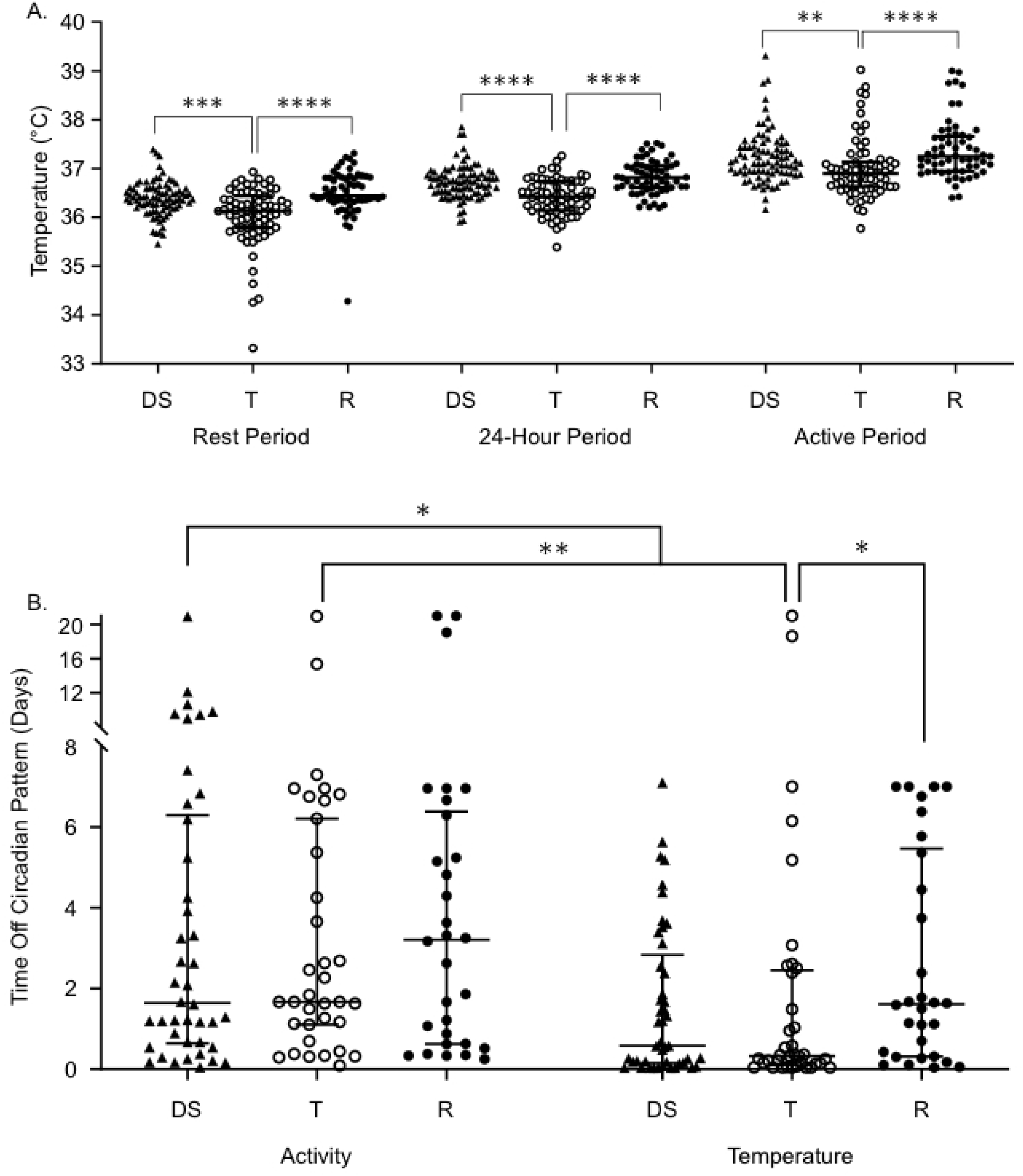
Tolerant mice have cooler body temperatures and deviate from normal body temperature patterns sooner than resistant mice. A. Core body temperature during resting, 24-hour, and active periods for delayed susceptible (DS, filled triangles), tolerant (T, open circles), and resistant (R, filled circles) mice. B. Time to deviation from circadian pattern (“off pattern”) for temperature and activity. Each symbol represents an individual mouse. Median and interquartile range indicated. Kruskal-Wallis test was performed to find significant differences (* P <0.05, ** P <0.01, *** P <0.001, **** P <0.0001).

As body temperature can be altered by differences in activity, activity levels were also analyzed for the fraction of time the mouse was active throughout a given time period. The fraction of time the mouse was active for their rest period, active period, and an entire 24-hour period was not significantly different between these groups (Fig S2).

Gross alterations in circadian patterns of core body temperature and locomotor activity occurred in nearly all infected lines of CC mice, but with varying times of onset and varying severity. STm tolerant mice deviated from their normal pattern of core body temperature very quickly, with a median of 463 minutes (7.7 hours post-infection) (Fig 3B), and without showing other perceptible clinical signs of disease. Delayed susceptible strains also deviated from their normal pattern of body temperature rapidly (835 minutes, ∼13 hours). Resistant mice maintained their circadian pattern significantly longer than tolerant mice with a median of 2,324.5 minutes (1.61 days) (Fig 3B, *P* = 0.0458), but despite their resistance to infection, they had perceptible changes in the regulation of their core body temperature.

All groups maintained their circadian pattern of locomotor activity for a similar amount of time – between 1 and 3 days post-infection (Fig 3B). When the time to deviation from activity and temperature patterns (time to “off-pattern”) were compared for each group, both delayed susceptible and tolerant mice had disruptions in their temperature pattern sooner than disruptions in their activity patterns, suggesting that one does not immediately influence the other (Fig 3B). Delayed susceptible mice deviated from their temperature pattern a median of 1,523.5 minutes (1.06 days) earlier than their activity pattern (Fig 3B, *P* = 0.0351), while tolerant mice deviated from their temperature pattern a median of 1,940 minutes (1.34 days) earlier (Fig 3B, *P* = 0.0031). Disrupted circadian patterns of temperature and activity were not significantly different in resistant mice (Fig 3B, *P* = >0.9999). For delayed susceptible and tolerant mice, changes in core body temperature pattern signaled earlier changes in health than did changes in activity level, consistent with our previous reports [37].

### Tolerant strains have increased total white blood cell counts, driven by circulating neutrophils, relative to resistant strains

Complete blood counts (CBC) were performed pre-infection and at the time of necropsy for mice that were euthanized at one week or three weeks post-infection [37]. The difference between pre-infection and post-infection CBC components was calculated to determine the change in each component (Table S2). We included only tolerant and resistant strains in this analysis because they were euthanized at the same time point. At three weeks post-infection, tolerant mice had significantly more circulating white blood cells (WBC) than resistant mice (5.29 x10^9^/L vs. 2.61 x10^9^/L, Fig 4A, *P* = 0.0396). The differential count suggested that circulating neutrophils (NEU) were elevated in tolerant mice relative to resistant mice at three weeks post-infection (Fig 4B, 5.58 x10^9^/L vs. 2.16 x10^9^/L, *P* = 0.0029). Circulating monocyte (MON) and lymphocyte (LYM) numbers were similar between tolerant and resistant mice (Fig 4C and 4D, *P* = 0.2278, >0.9999). Thus, circulating neutrophils in tolerant mice remain elevated for the duration of the study period.

**Figure 4:**
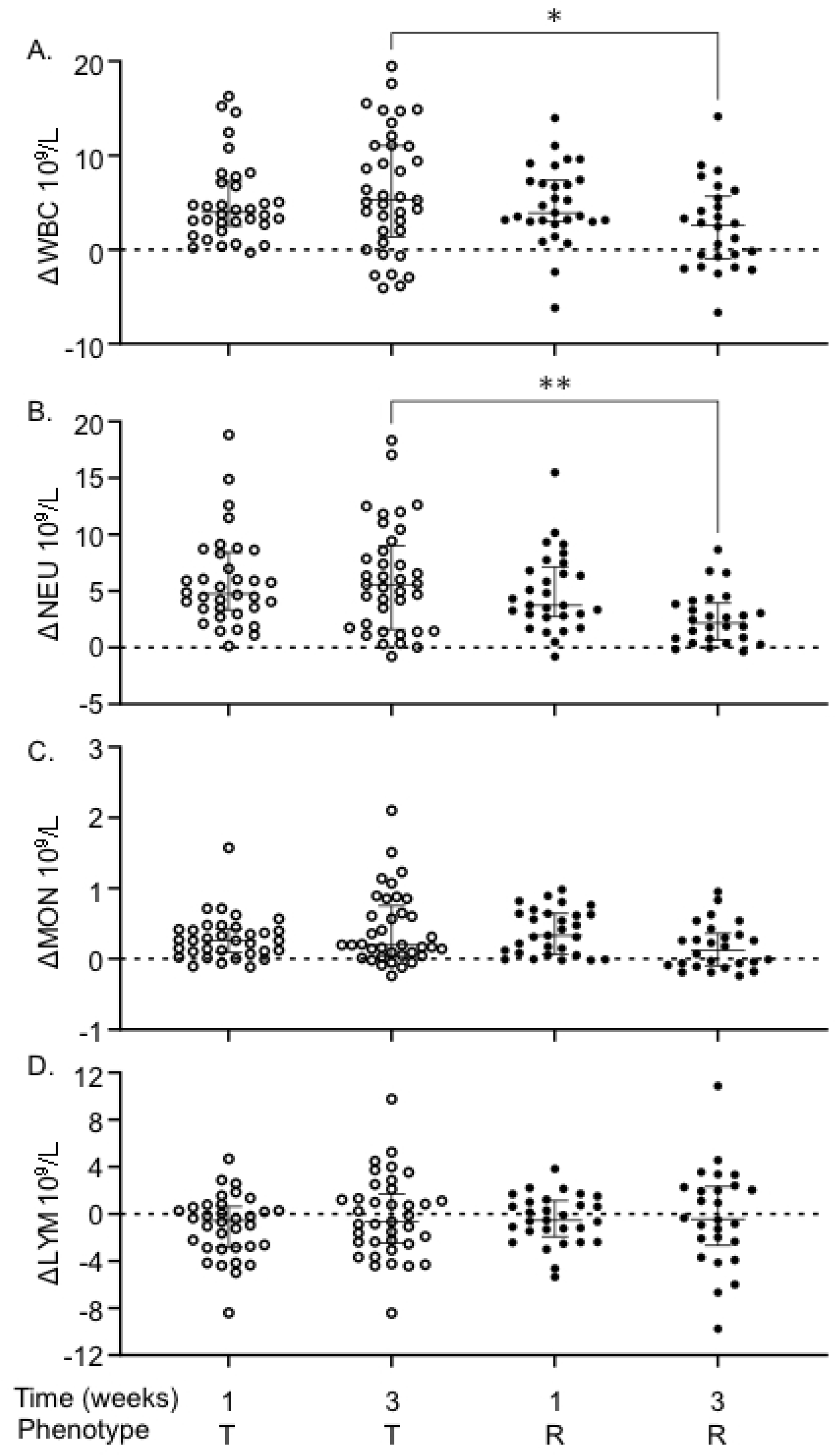
Tolerant mice have higher WBC and higher NEU than resistant mice. Differences between infected and uninfected for circulating A. total white blood cells (WBC), B. neutrophils (NEU), C. monocytes (MON), and D. lymphocytes (LYM). Each circle represents an individual mouse and strains represent medians and interquartile ranges. T = tolerant (open circles), R = resistant (filled circles). Kruskal-Wallis test was performed to find significant differences (* *P* <0.05, ** *P* <0.01).

### At three weeks post-infection, tolerant mice have increased MCP-1 and IFN-γ levels and reduced ENA-78 levels compared to resistant mice

Chemokines, cytokines, and their receptors control the movement of cells of the immune system. The levels of 36 cytokines/chemokines in serum from uninfected, one week, and three weeks post-infection CC mice were determined. The change in circulating cytokine/chemokine levels between uninfected and infected animals was calculated (Table S3). We included only tolerant and resistant strains in this analysis because they were euthanized at the same time point, while delayed susceptible animals were euthanized earlier. STm tolerant and resistant mice had different levels of three cytokines at three weeks post-infection: MCP-1, IFN-γ, and ENA-78.

MCP-1 (CCL2) is a key chemoattractant for monocytes and plays a critical role in their migration and tissue infiltration [38]. The change in serum MCP-1 over baseline is similar for tolerant and resistant lines at one week post-infection. However, at three weeks post-infection, MCP-1 levels remain elevated in tolerant mice in response to infection, a median increase of 117.54 pg/ml, while circulating MCP-1 returns nearly to baseline in resistant mice (increase of 9.27 pg/ml over baseline (Fig 5A, *P* = 0.0005)). Serum MCP-1 fell in resistant mice from a median increase of 123.57 pg/ml over baseline at one week post-infection to a median increase of 9.27 pg/ml at three weeks post-infection (Fig 5A, *P* = <0.0001).

**Figure 5:**
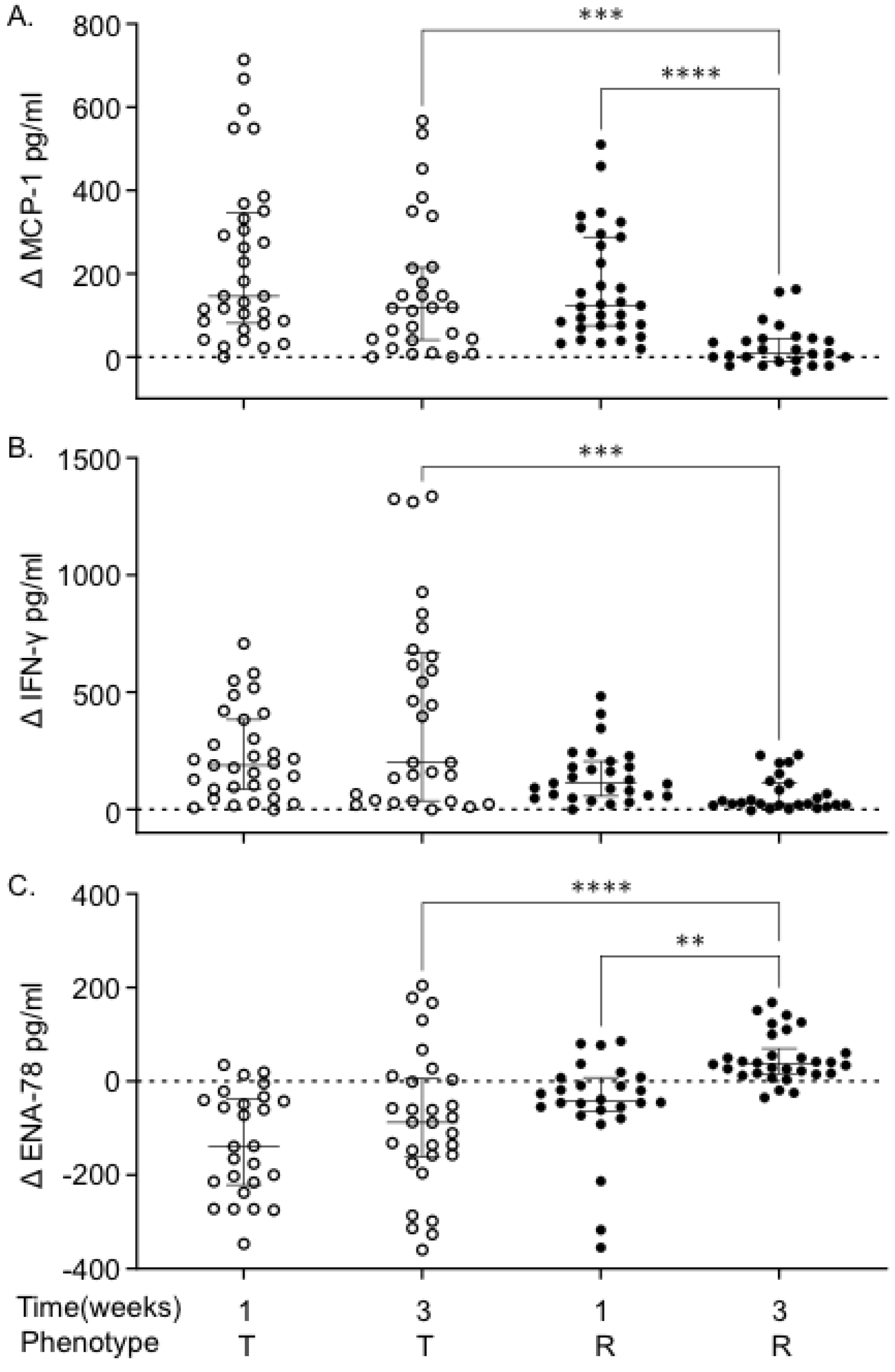
Tolerant strains have higher circulating MCP-1 and INF-γ, but lower ENA-78 at 3 weeks post-infection than resistant strains. The differences between infected and uninfected levels of, A. MCP-1, B. IFN-γ, and C. ENA-78 in tolerant and resistant strains. Circles represent individual mice and strains represent medians and interquartile ranges. Outliers were removed. T = tolerant (open circles), R = resistant (closed circles). Kruskal-Wallis test was performed to find significant differences (* *P* <0.05, ** *P* <0.01, *** *P* <0.001, **** *P* <0.0001).

IFN-γ is vital for controlling intracellular replication of STm, promoting damage of the *Salmonella* containing vacuole (SCV), the intracellular niche of STm, and promoting production of reactive oxygen and nitrogen species from phagocytic cells [39]. At one week post-infection, IFN-γ levels rose similarly in tolerant versus resistant mice. However, tolerant strains maintained a high level of circulating IFN-γ with a median increase of 200.04 pg/ml over baseline at three weeks post-infection, while resistant strains returned IFN-γ levels much closer to baseline (median increase of 25.99 pg/ml (Fig 5B, *P* = 0.0001)).

ENA-78 (CXCL5), is a CXC motif chemokine normally produced by platelets that promotes accumulation of neutrophils [40]. Tolerant strains had significantly lower circulating ENA-78 levels than STm resistant strains at three weeks post-infection compared to the baseline (Fig 5C, −87.90 pg/ml vs. 37.91 pg/ml, *P* = <0.0001). Resistant strains also significantly increased their ENA-78 levels from one week to three weeks post infection from −42.11 pg/ml to 37.91 pg/ml over uninfected baseline (Fig 5C, *P* = 0.0016). Thus, reduction of circulating CXCL5 appears to improve host defense to bacterial infection perhaps by allowing proper scavenging of other chemokines (keratinocytes-derived chemokine (KC or CXCL1) and macrophage inflammatory protein 2-alpha (MIP2-α or CXCL2)), allowing the formation of chemokine gradients and sensitizing CXCR2 resulting in an increased influx of neutrophils [41].

### Tissue damage is more severe in tolerant and delayed susceptible strains than in resistant strains

Sections of ileum, cecum, colon, spleen, and liver were stained with H&E and were scored blindly by a board-certified pathologist using the same scoring system as previously described [37] of 0 to 4 (0 = normal, 4 = severe damage, Table S4). Intestinal organs had minimal damage (scores 0-1) and were excluded from further analysis (Fig S3). Spleen and liver had a range of damage and were examined further at one and three weeks post-infection in tolerant and resistant strains only.

When mice were grouped by response to infection, there was no significant difference between delayed susceptible, tolerant, and resistant for spleen or liver damage at one week post-infection. However, strains susceptible to acute infection had significantly more splenic damage at 7 days post-infection than strains of other phenotypes (Fig S4). At three weeks post-infection, however, significant differences became apparent. STm tolerant strains had a mean histopathology score of 1.89 (± 1.43), while delayed susceptible strains had a mean of 2.93 (± 1.47) for spleen, damage to these tissues was not statistically significantly different (Fig 6). Resistant strains had a mean of 0.53 (± 0.77) for spleen, lower than for both tolerant and delayed susceptible strains (Fig 6, *P* = 0.0004, <0.0001). Delayed susceptible strains also had increased spleen damage from 1.68 (± 1.26) at one week post-infection to 2.933 (± 1.47) at three weeks post-infection (Fig 6, *P* = 0.007).

**Figure 6:**
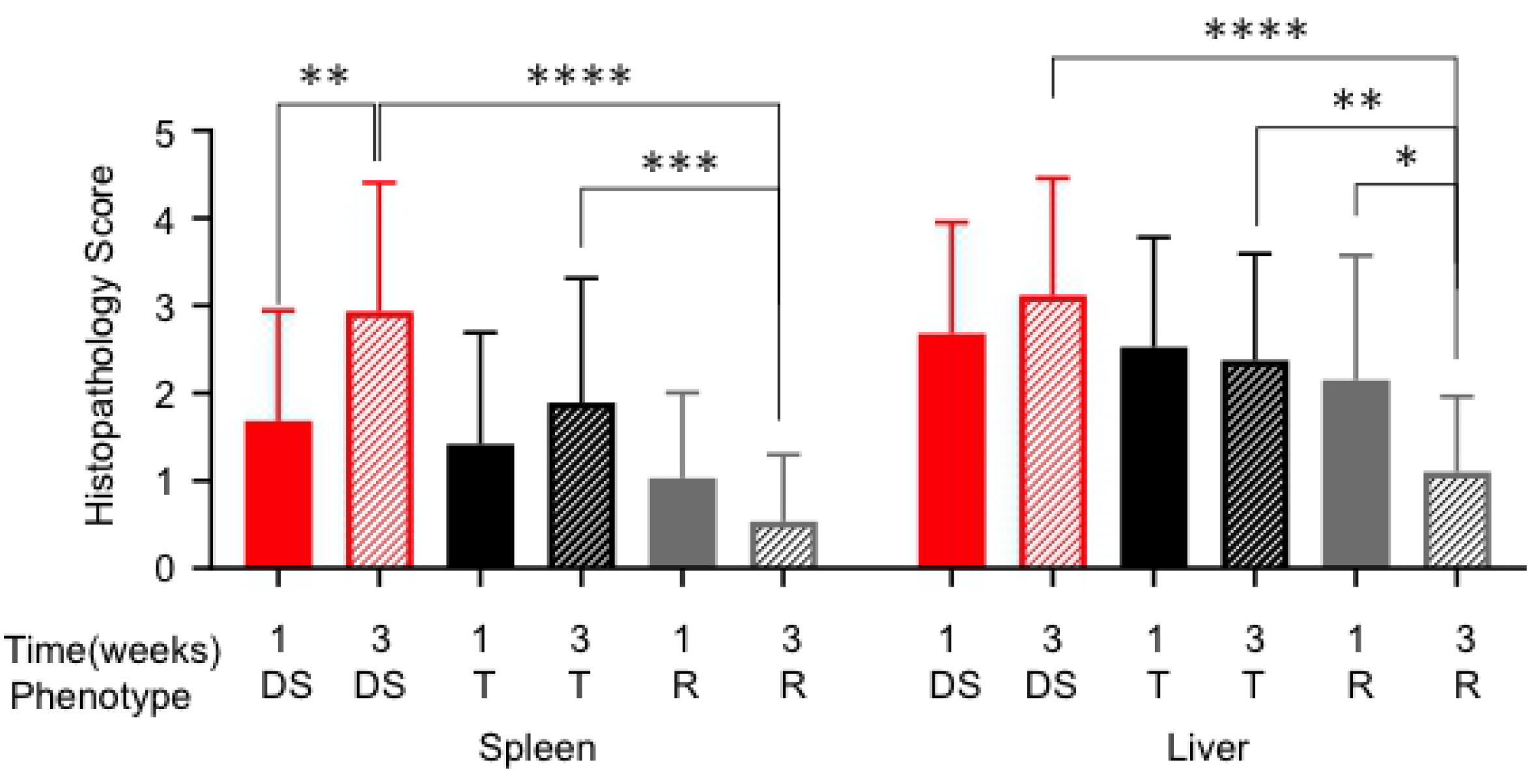
Resistant strains have less damage in spleen and liver at three weeks post-infection than tolerant and delayed susceptible strains. The mean of spleen and liver histopathology scores for tissue damage and standard deviation indicated. T = tolerant (black bars), R = resistant (gray bars), DS = delayed susceptible (red bars). Kruskal-Wallis test was performed to find significant differences (* *P* <0.05, ** *P* <0.01, *** *P* <0.001, **** *P* <0.0001).

Although damage to the liver was more pronounced than splenic damage at three weeks post-infection, the trend was the same: tolerant strains suffered more severe damage than resistant strains (tolerant mean of 2.38 (± 1.22) and resistant mean of 1.10 (± 0.87)) (Fig 6, *P* = 0.0058). Delayed susceptible strains also suffered more liver damage at three weeks post-infection than resistant strains (delayed susceptible mean of 3.12 (± 1.34) and resistant mean of 1.10 (± 0.87)) (Fig 6, *P* = <0.0001). Liver damage was scored as resolving between one and three weeks post-infection in resistant strains (mean of 2.15 (± 1.42) at one week to 1.10 (± 0.87) at three weeks) (Fig 6, *P* = 0.0337).

### Significant and suggestive genetic associations were identified across various phenotypes for three weeks post-STm infection

All 18 CC strains infected here were included in a quantitative trait loci (QTL) association of relevant phenotypes. A statistically significant association was identified for spleen colonization on mouse Chromosome (Chr) 3 (Spleen colonization QTL 1 (*Scq1)*) (Fig 7 and Table 1). QTL *Scq1* contains 533 genes, with no obvious haplotype differences, likely due to the small number of strains in our analysis (Fig 7A and 7B). A suggestive association was identified for liver colonization on Chr 13, containing 20 genes. The NZO founder has a high haplotype effect in this region, associated with an increase in liver colonization, and 3 CC strains carried this haplotype (CC001 (DS), CC003 (DS), and CC072 (T)) (Fig 7C and 7D). Furthermore, the B6 founder strain has a low haplotype effect in this region, associated with reduced liver colonization, and 3 CC strains carried this haplotype (All resistant: CC015, CC057, CC058) (Fig 7C and 7D). Of the 20 genes in this region on Chr 13, none had SNP differences matching the haplotype effects.

**Figure 7:**
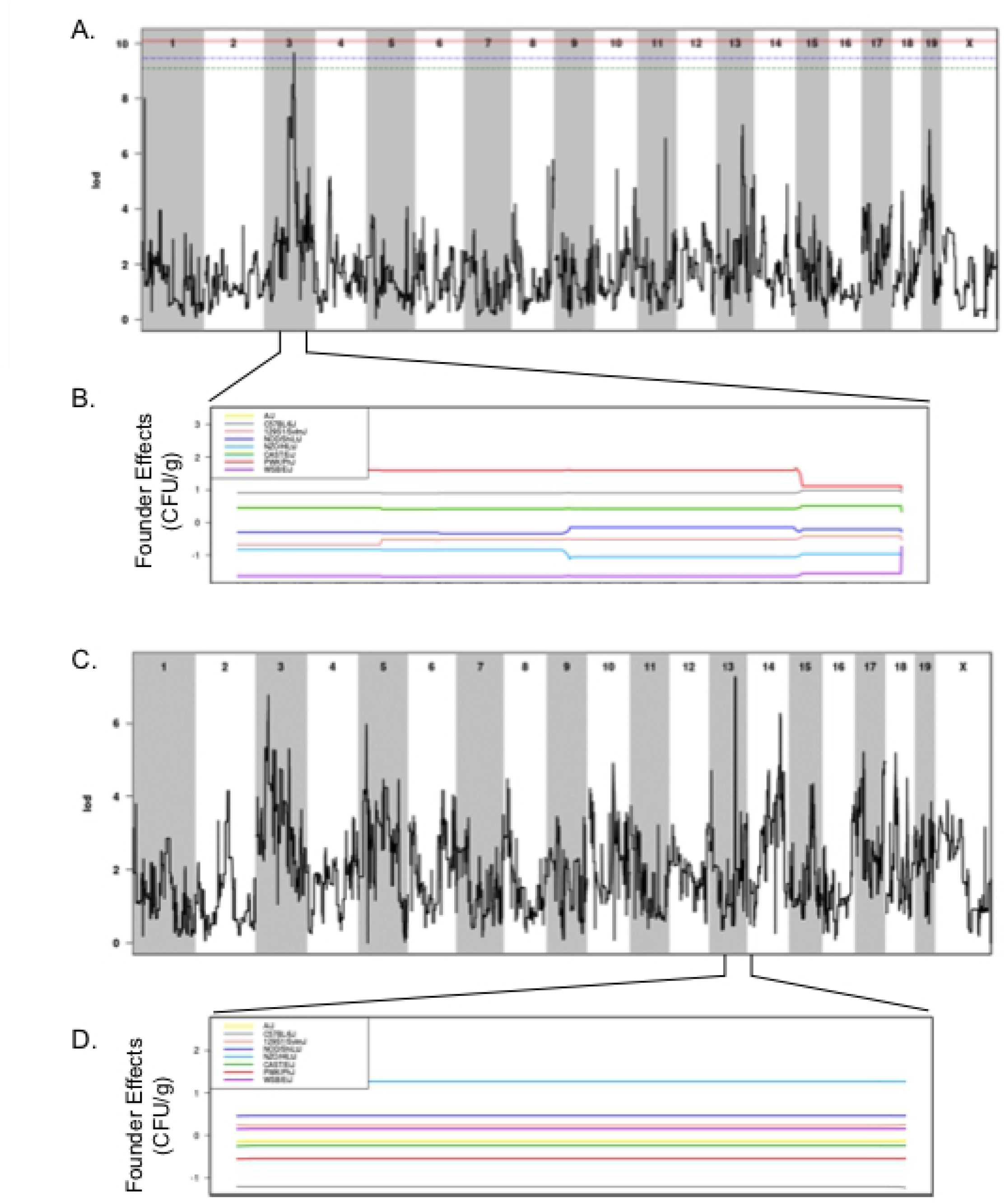
QTLs identified for spleen and liver colonization using 18 CC strains. A. QTL of spleen CFU at three weeks post-infection and B. allele effect plots for Chr 3. C. QTL of liver CFU at three weeks post-infection and D. allele effect plots focused in on Chr 13. Green dotted line is 85% significant, blue dotted line is 90% significant, red dotted line is 95% significant. Results obtained using gQTL.

**Table 1:** QTL associations for spleen and liver colonization and resistance and tolerance categorization after STm infections. 18 CC strains were included in spleen and liver colonization QTL, 32 CC strains were in the resistance QTL, and 27 CC strains were in the tolerance QTL.

| QTL name | Phenotype | Chr | LOD | P-value | Proximal (Mb) | Max (Mb) | Distal (Mb) | Haplotype Effect |
| --- | --- | --- | --- | --- | --- | --- | --- | --- |
| | Resistance | 3 | 8.44 | $2.07 \times 10^{-6}$ | 95.34 | 95.94 | 100.14 | high: NZO, 129S1, WSB |
| | Resistance | 13 | 9.37 | $3.10 \times 10^{-7}$ | 81.53 | 81.87 | 83.81 | high: B6, PWK |
| | Resistance | 17 | 6.85 | $4.88 \times 10^{-5}$ | 86.91 | 88.97 | 90.81 | high: PWK, NZO, CAST |
| | Tolerance | 2 | 7.9 | $6.19 \times 10^{-6}$ | 159.05 | 168.27 | 181.43 | high NOD, NZO |
| | Tolerance | 6 | 6.53 | $9.14 \times 10^{-5}$ | 114.37 | 147.33 | 147.37 | |
| *Scq1* | Spleen Colonization | 3 | 9.64 | $1.78 \times 10^{-7}$ | 79.59 | 93.37 | 95.95 | |
| | Liver Colonization | 13 | 7.28 | $2.11 \times 10^{-5}$ | 81.83 | 83.66 | 83.8 | high: NZO, low: B6 |

To overcome the limitations of using small numbers of CC strains to explore the genes that distinguish tolerant and resistant strains, all 32 CC strains whose acute response to infection with STm is known were included in a binary QTL analysis [37]. Resistant strains (CC015, CC024, CC051, CC057, CC058) were assigned a value of one while the remaining tolerant, delayed susceptible, and susceptible strains were assigned a value of zero. This analysis identified three suggestive QTL on Chr 3, 13, and 17 (Fig 8 and Table 1). The Chr 3 QTL contains 212 genes and high haplotype effects in NZO (All resistant: CC015, CC058), 129S1 (CC024 (R), CC037 (S), CC053 (DS), CC057 (R)), and WSB (CC051 (R)) suggested that regions from these founders conferred resistance (Fig 8A and 8B). No SNP differences were identified in this region between NZO, 129S1, and WSB and the remaining CC founder strains.

**Figure 8:**
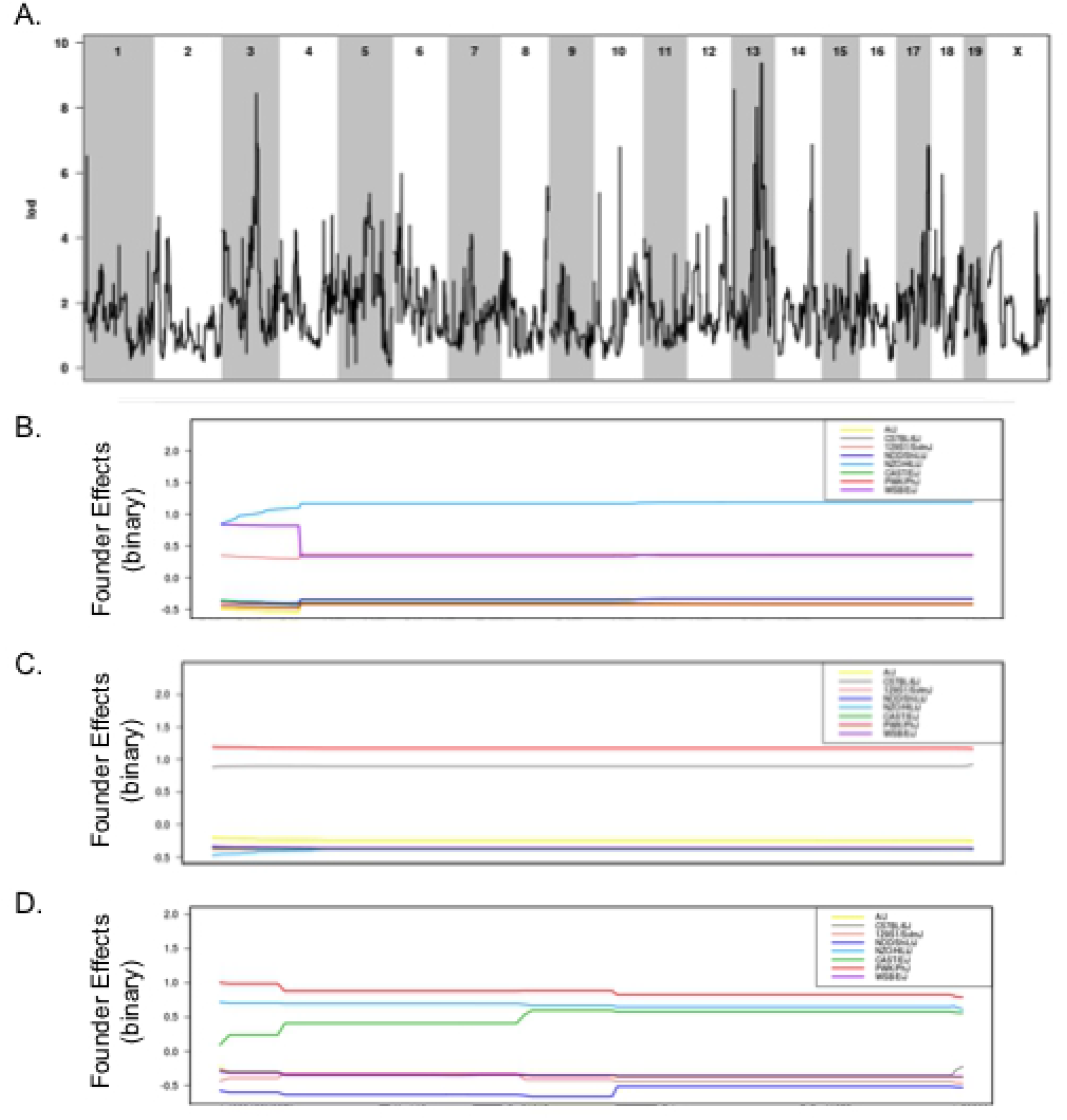
Binary categorization to identify QTLs linked to STm resistance using 32 CC strains. A. QTL of resistant categorization (score of 1) versus susceptible, delayed susceptible, and tolerant categorization (score of 0) after STm infections. Allele effect plots zoomed in on B. Chr 3, C. Chr 13, and D. Chr 17. Results obtained using gQTL.

The association on Chr 13 contained 29 genes and had a high haplotype effect (more STm resistant) for PWK (CC024 (R)) and B6 (CC015 (R), CC030 (S), CC057 (R), CC058 (R)) (Fig 8A and 8C). There were no genes in this region that had SNP differences that followed the haplotype effects. The association on Chr 17 contained 64 genes and had a high haplotype effect (more resistant to STm) for PWK (CC024 (R)), NZO (CC013 (S), CC051 (R), CC057 (R)), and CAST (CC015 (R), CC019 (S), CC058 (R)) (Fig 8A and 8D). Only one gene had a SNP difference that corresponded to the haplotype effects.

Tolerance to STm infection was also examined by assigning tolerant strains (CC002, CC017, CC038, CC043, CC072, CC078) a score of one and susceptible and delayed susceptible strains a score of zero. A total of 27 CC strains were included in this analysis (resistant strains were excluded). This analysis yielded suggestive associations with QTL on Chr 2 and Chr 6 (Fig 9 and Table 1). The associated region on Chr 2 contains 618 genes, and NOD (All tolerant: CC072, CC078) and NZO (All tolerant: CC002, CC043) had a high haplotype effect. Thus, strains that had a NOD or NZO allele at this location were more likely to be tolerant to STm infections (Fig 9A and 9B). There were no genes in the Chr 2 region that had SNP differences corresponding to haplotype effects. The association on Chr 6 had 751 genes, but no haplotype effects were identified (Fig 9A and 9C). While the experiments examining spleen and liver bacterial burdens did not contain enough strains to identify significant associations, the binary categorizations did not capture enough phenotypic diversity to be significant.

**Figure 9:**
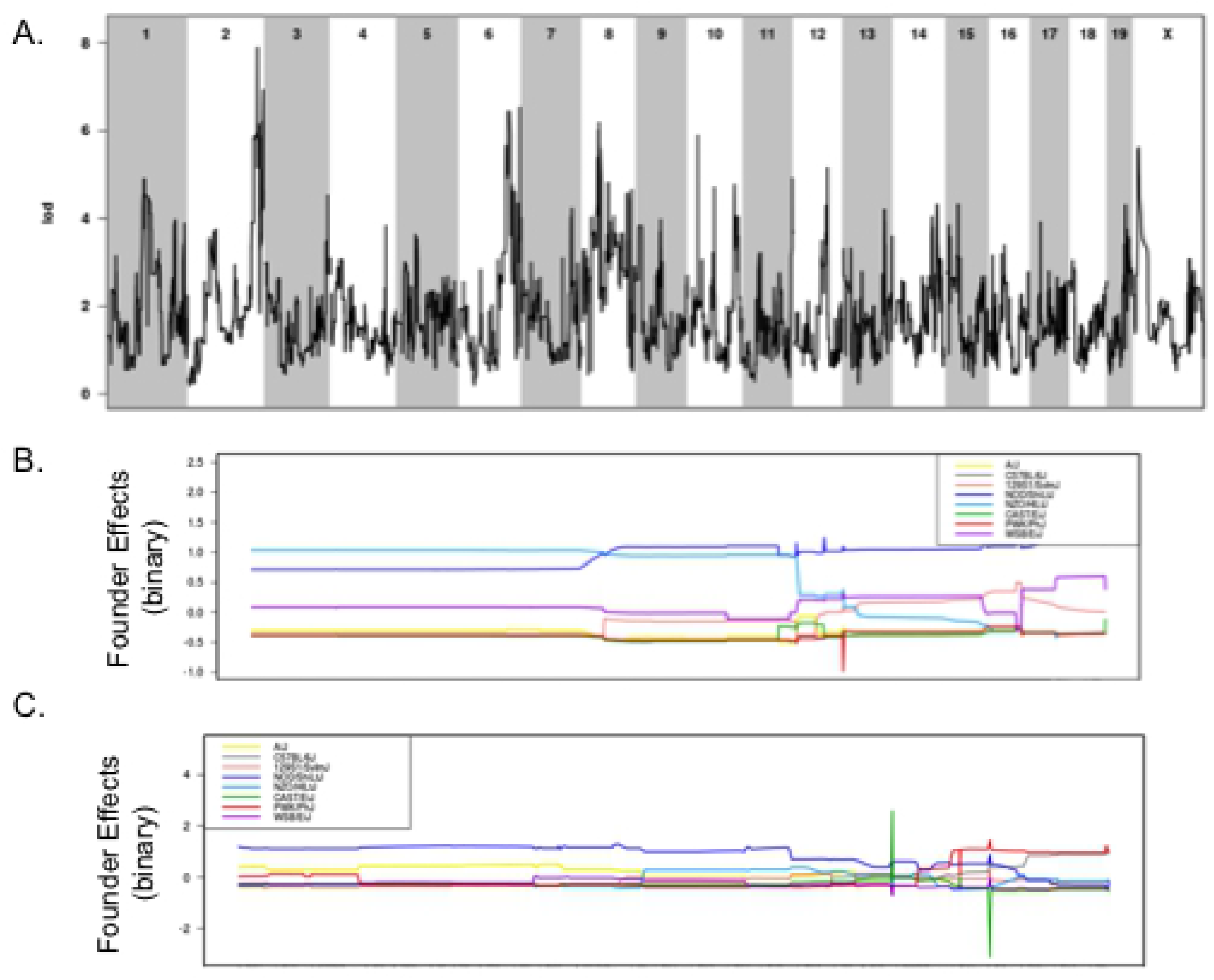
Binary categorization to identify QTLs linked to STm tolerance using 27 CC strains. A. QTL of tolerant categorization (score of 1) versus susceptible and delayed susceptible categorization (score of 0) after STm infections. Allele effect plots zoomed in on B. Chr 2 and C. Chr 6. Results obtained using gQTL.

### Transcriptional differences in QTL regions in tolerant versus resistant mice

RNA sequencing on spleen samples of uninfected and infected mice identified differentially expressed genes. Samples from infected animals were normalized to strain-specific samples from uninfected animals and then tolerant and resistant group means were compared to identify differential expression across these phenotypes. Data were normalized to the tolerant group, so a positive value indicates up-regulation of a given gene in the resistant group and a negative value indicates up-regulation in the tolerant group. Only genes that were at least 2-fold differently expressed, and statistically significantly changed (*P* = <0.05), were included (552 genes). These genes were mapped to the QTL regions we identified in the previous section.

*Scq1*, the QTL region linked to STm spleen colonization that we identified on Chr 3, contains 533 genes. Five genes from this region were differentially expressed: fibrinogen alpha chain (*Fga*), fibrinogen beta chain (*Fgb*), fibrinogen gamma chain (*Fgg*), S100 calcium binding protein A5 (*S100a5*), and trichohyalin (*Tchh*) (Table 2). *Fga*, *Fgb*, *Fgg*, and *Tchh* were all up-regulated in resistant strains, while *S100a5* was up-regulated in tolerant strains. *Fga*, *Fgb*, and *Fgg* are the three subunits of fibrinogen and are involved in clot formation, wound healing, and microbial resistance and are thus strong candidates for further study [42]. *S100a5* is expressed primarily in neuronal bodies [43] and *Tchh* is expressed in the tongue and hair [44], and do not appear to be involved in response to infection. Of the 20 genes found in the QTL linked to liver colonization on Chr 13, none were differentially expressed in the spleens of tolerant versus resistant mice.

**Table 2:**
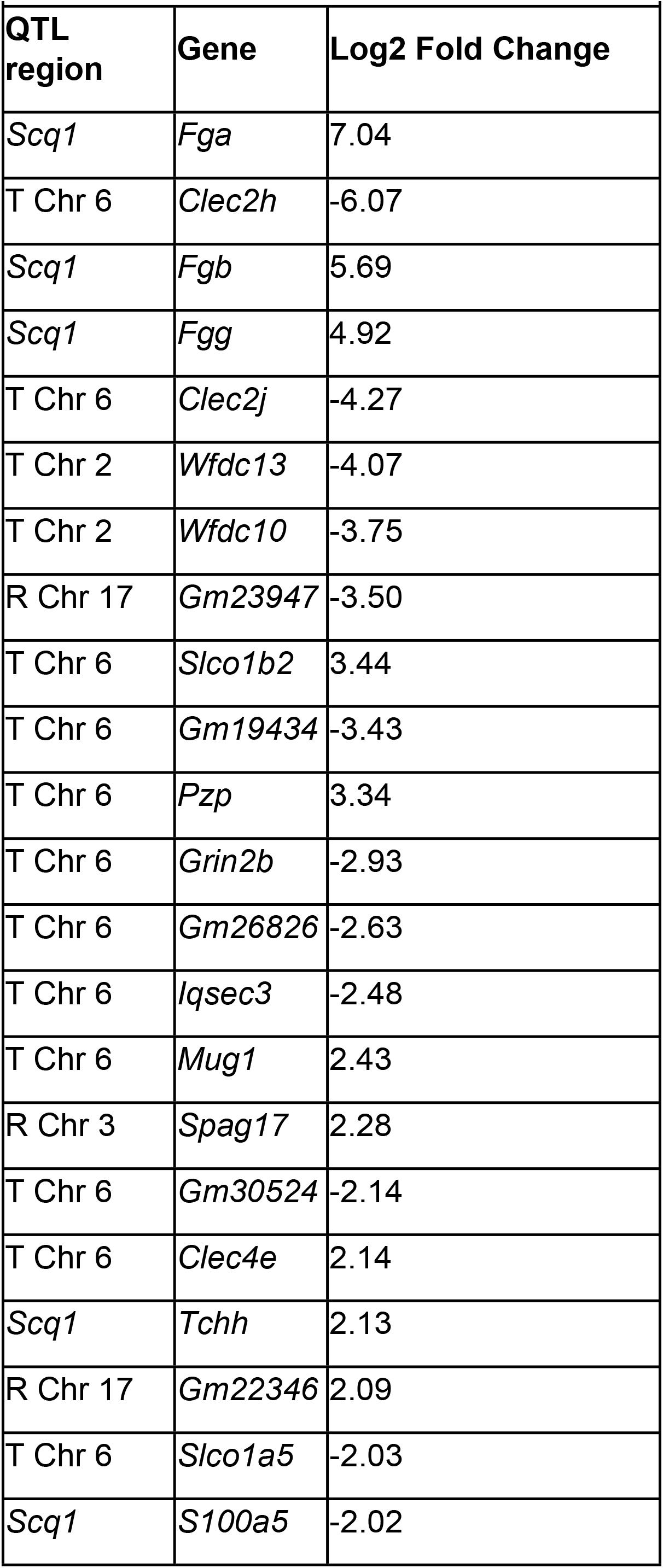
QTL associations that contain differentially expressed genes in STm tolerant vs. resistant CC mice. Tolerance was used as the baseline in RNA comparison, so log_2_ fold change values that are positive = up-regulated in resistant and negative values = up-regulated in tolerant.

One gene of the 212 genes in the QTL region on Chr 3 associated with spleen colonization (sperm-associated antigen 17 (*Spag17*)) and two predicted genes of the 64 genes in the QTL region on Chr 17 associated with liver colonization (*Gm22346* and *Gm23947*) were differentially expressed for the resistance phenotype (Table 2). *Spag17* and *Gm22346* were up-regulated in resistant strains, while *Gm23947* was up-regulated in tolerant strains. *Spag17* is involved in motility of axonemes, such as sperm flagella and the cilia of lung epithelium [45]. The two predicted genes linked to resistance are of unknown function. The QTL linked to resistance to STm infection on Chr 13 overlaps with the QTL associated with liver colonization on Chr 13 and does not contain any differentially expressed genes.

Of the genes located in the suggestive QTL on Chr 2 associated with the tolerance phenotype, two genes were differentially expressed: WAP four-disulfide core domain 10 (*Wfdc10*) and WAP four-disulfide core domain 13 (*Wfdc13*) (Table 2). These genes have antimicrobial functions and are up-regulated in response to exposure to LPS [46]. On the suggestive QTL on Chr 6 associated with tolerance, three predicted genes and nine genes were differentially expressed (Table 2). C-type lectin domain family 2, member h (*Clec2h*), C-type lectin domain family 2, member J (*Clec2j*), and C-type lectin domain family 4, member e (*Clec4e*) are members of the same family and are involved in immunity. *Clec2h* and *Clec2j* are up-regulated in tolerant strains, while *Clec4e* is up-regulated in resistant strains. *Clec2h* is expressed in the intestines and is involved in intestinal monitoring [47, 48], while *Clec4e* is involved in macrophage functioning as well as cytokine induction [49]. *Clec2j* function is unknown [50]. PZP, alpha-2-macroglobulin like (*Pzp*) is an immunosuppressive protein often expressed during pregnancy and high levels are linked with poor outcomes to infection [51, 52]. *Pzp* is up-regulated in resistant strains. Solute carrier organic anion transporter family, members 1a5 (*Slco1a5*) and 1b2 (*Slco1b2*) are in the same family. *Slco1a5* and *Slco1b2* are involved in bile acid and bile salt transport and deletion of them leads to a buildup of bilirubin in the blood [53, 54]. These genes play a key role in drug uptake, specifically in the liver and intestines. *Slco1a5* is up-regulated in tolerant strains, while *Slco1b2* is up-regulated in resistant strains.

### Ingenuity pathway analysis revealed pathways involved in response to infection

RNA sequencing data from both uninfected and infected spleen samples identified putative pathways involved in the response to STm infection. Ingenuity pathway analysis (IPA) identified the top canonical pathways (Table 3).

**Table 3:** Top canonical pathways for spleen RNA-seq comparing tolerant and resistant mice at three weeks post-infection identified using Ingenuity Pathway Analysis. Top 5 pathways listed. Genes found in QTL regions highlighted in red.

| Table 3: IPA Top 5 Canonical Pathways |  |  |  |  |
| --- | --- | --- | --- | --- |
| Ingenuity Canonical Pathways | -log (p-value) | Ratio | z-score | Molecules |
| Acute Phase Response Signaling | 15.00 | 0.11 | 1.39 | ALB, AMBP, APCS, APOA1, APOA2, APOH, C9, CRP, FGA, FGB, FGG, HPX, IL36G, MBL2, PLG, SAA1, SAA2-SAA4, Saa3, SERPINA3, TTR |
| LXR/RXR Activation | 14.70 | 0.14 | 2.67 | ALB, AMBP, APOA1, APOA2, APOB, APOH, C9, FGA, GC, HPX, IL1R2, IL1RAPL1, IL36G, KNG1, NOS2, SAA1, TTR |
| FXR/RXR Activation | 13.20 | 0.13 | NaN | ALB, AMBP, APOA1, APOA2, APOB, APOH, C9, FBP1, FGA, GC, HPX, IL36G, KNG1, SAA1, SLCO1B3, TTR |
| Xenobiotic Metabolism PXR Signaling Pathway | 7.43 | 0.07 | 1.94 | CES3, CES5A, CHST4, CYP2B6, CYP2C8, CYP3A5, ESD, NOS2, SULT4A1, UGT1A9 (includes others), UGT2B17, UGT2B28, UGT2B7 |
| LPS/IL-1 Mediated Inhibition of RXR Function | 6.05 | 0.05 | -1.13 | CHST4, Cyp2a12/Cyp2a22, CYP2B6, CYP2C8, Cyp2d9 (includes others), Cyp2j5, CYP3A5, IL1R2, IL1RAPL1, IL36G, SLCO1A2, SLCO1B3, SULT4A1 |

The five pathways with the highest log score were Acute Phase Response Signaling, LXR/RXR Activation, FXR/RXR Activation, Xenobiotic Metabolism PXR Signaling Pathway, and LPS/IL-1 Mediated Inhibition of RXR Function (Table 3). Acute Phase Response Signaling, LXR/RXR Activation, and Xenobiotic Metabolism PXR Signaling Pathway were up-regulated in resistant mice. LPS/IL-1 Mediated Inhibition of RXR Function was up-regulated in tolerant mice. Acute Phase Response Signaling, LXR/RXR Activation, and FXR/RXR Activation all contained the gene *Fga* and Acute Phase Response Signaling contained the genes *Fgb* and *Fgg*. *Fga*, *Fgb*, and *Fgg* were also found in the QTL regions associated with spleen colonization, suggesting that these genes are the most likely to differentiate between resistance and tolerance.

## Discussion

A variety of host immunologic, physiologic and metabolic factors likely contribute to differences in disease outcome after exposure to an infectious agent. In previous work, we described our observations around susceptibility and survival after acute *S*. Typhimurium infection in genetically diverse mice [37]. However, we found that infection periods longer than 7 days were required to differentiate tolerant and resistant phenotypes because bacterial burden and survival were similar between these strains at one week post-infection [31,32,37]. To differentiate these host disease phenotypes, we infected 18 strains of CC mice, known to survive acute infection, with STm for up to three weeks [37].

Seven of these 18 CC strains did not survive three week post-infection and thus had a survival phenotype we called “delayed susceptible.” Six of the remaining strains fit the criteria for tolerance, surviving infection with a bacterial load that caused serious systemic infection in delayed susceptible strains. Resistant strains were colonized at very low levels compared to their tolerant counterparts and exhibited no clinical signs of disease. When all resistant strains were grouped, they had a median colonization of 1.26 x10^2^ CFU/g in liver and 2.82 x10^3^ CFU/g in spleen at three weeks, significantly less than tolerant strains that had a median of 1.29 x10^4^ CFU/g in liver and 1.13 x10^5^ CFU/g in spleen (Fig S6, both *P* = <0.0001).

PP and MLN colonization varied across the CC strains we examined. Long-term STm colonization in mice is linked to the persistence of infection in the RE system, particularly in the MLN [55]. In our study, tolerant strains, defined by persistently higher bacterial load, had significantly more STm in their MLN than resistant strains, supporting a role for the RE system in long-term colonization.

We previously showed that pre-infection baseline core body temperature differs between strains that survived or were susceptible to acute STm infections. Surviving strains had cooler pre-infection baseline temperatures by ∼0.3 °C across their resting period, active period, and full 24-hour period [37]. This difference did not appear to be related to higher or lower locomotor activity [37], but could be the result of differences in metabolism. In our current work, tolerant strains had the lowest baseline core body temperatures across all three time periods compared to resistant and delayed susceptible strains. Tolerant strains also had cooler body temperatures when compared to strains susceptible to acute infection by ∼0.5 °C.

Disruption of circadian patterns of body temperature and activity have previously been reported during parasitic and viral infections [56, 57]. We recently reported that circadian patterns of body temperature and activity are disrupted in CC mice acutely infected with STm [37], and this was also true during the longer post-infection periods described here. After infection, the time to disrupted circadian patterns for core body temperature was different between resistant and tolerant strains. Diurnal patterns of core body temperature were disrupted earlier in strains that were tolerant of STm infection than in resistant strains. Thus, in tolerant strains that survive STm infection despite showing signs of disease, we can see earlier symptoms with telemetry that are not otherwise readily observable [57, 58]. Lipopolysaccharide (LPS) injection in rats is correlated with suppression of biological clock genes [59]. Although LPS may contribute to the circadian pattern disruption in our experiments, different onset and timing of pattern disruption in mice with different genetic backgrounds suggest that this disruption is more complex than previously appreciated.

Circadian patterns of locomotor activity in infected mice were disrupted later than patterns of core body temperature in all groups of mice (Fig 3B), although the time to disruption did not vary between STm tolerant and resistant CC mice. Early disruption of body temperature rhythms versus other clinical signs has also been observed in SIV infected monkeys [57]. Detection of circadian pattern disruption was also compared between temperature and activity for each phenotypic group. Disruption of temperature patterns was a more sensitive predictor of disease onset for tolerant and delayed susceptible strains, but not for resistant strains.

Differences in the immune response were observable between tolerant and resistant strains. Tolerant strains had a larger rise in white blood cell count driven by a significantly higher increase in neutrophils than resistant strains after infection with STm. Neutrophils are part of the innate immune system and are vital to early control of infections [10]. Tolerant animals appear to maintain neutrophilia secondary to prolonged infection with STm. Since resistant strains appear to be clearing the infection, the lower bacterial count may reduce the need for large numbers of neutrophils. Histopathology showed large numbers of neutrophil infiltration in both spleen and liver primarily in tolerant mice, supporting this.

We discovered 3 cytokines that were significantly different between tolerant and resistant strains. IFN-γ was increased in both resistant and tolerant strains, but tolerant strains had a much larger increase after infection. IFN-γ keeps STm at a manageable level for the host, and neutralizing IFN-γ results in more rapid replication of STm increasing the severity of infection [55]. Since tolerant strains have such a high level of IFN-γ, this likely contributes to controlling their bacterial burden and perhaps their disease symptoms.

MCP-1 (CCL-2) is also elevated in STm tolerant mice relative to their resistant counterparts. This CC chemokine is a chemoattractant for monocytes and macrophages but also for lymphocytes, NK cells, basophils, and dendritic cells [60], and is produced by many cell types. Septic human patients also have elevated serum MCP-1 [61], which can be induced during inflammation by cytokines and chemokines including IFN-γ and other substances like LPS [62]. High levels of MCP-1 have been linked to more severe organ damage/failure, higher levels of septic shock, and higher mortality [63, 64]. Transgenic mice that overexpress MCP-1 are more highly susceptible to infections with intracellular bacterial pathogens [62, 65]. In models with reduced serum MCP-1, these animals are resistant to infection with Listeria monocytogenes [65]. High serum levels of MCP-1 may blind cellular responses to local levels of MCP-1 [62, 65]. Finally, polymorphisms in the CCL2 gene have been shown to influence the outcomes of infection with Mycobacterium tuberculosis [66]. The MCP-1 elevation in our STm tolerant and resistant animals are consistent with these findings, yet the tolerant animals are able to survive the increased bacterial load that appears to occur with elevated MCP-1. Since STm resides and replicates in monocytes, high levels of MCP-1 may not be protective [67].

Finally, ENA-78, decreased in tolerant mice and increased in resistant mice, is a neutrophil chemoattractant [68]. Tolerant strains have a significantly higher number of circulating neutrophils than resistant strains, despite having less circulating ENA-78. Hepatocytes secrete ENA-78 in response to bacterial invasion or tissue damage [69]. Hepatocytes in tolerant strains may be creating the proper chemokine gradients that would attract an influx of neutrophils into the liver, while resistant strains that have started clearing bacteria and repairing damage no longer need an influx of neutrophils [41, 69]. Further experiments are required to confirm the role of this cytokine.

Tolerance to infectious agents has been hypothesized to correlate with reduced damage in infected tissues [33]. In our experiments, tolerant strains had milder tissue damage than strains that were susceptible to acute STm infection only in spleen at 7 days post-infection (Fig S4) [37]. However, strains that tolerated STm infection had equivalent tissue damage at one and three weeks post-infection, showing no evidence of repair (Fig 6). While resistant mice appeared to be actively repairing tissue damage by the three week time point. Furthermore, tolerant strains, which were colonized with STm at similar levels to delayed susceptible strains, had tissue damage that was as severe as tissue damage in DS strains. Tissue damage and repair have been linked to tolerance for other pathogens, yet we did not observe this [70, 71]. Our findings suggest that in our system, tissue damage and repair do not appear to distinguish tolerance from other phenotypes. Tolerance to STm infections likely relies on other physiologic, immunologic, and metabolic mechanisms [30, 71].

We used QTL analysis to identify genetic regions associated with resistance, tolerance, and in STm colonization of the spleen and liver. We identified one significant association with spleen colonization, multiple associations with resistance and tolerance, and one suggestive association with liver colonization. However, the small number of CC strains used and the lack of normal distribution of data did not allow us to pinpoint individual genes responsible for these associations [72]. RNA-seq on infected spleen allowed us to identify genes that are differentially expressed in tolerant versus resistant strains in response to STm infection, and to identify where these genes overlap with our QTL regions. Fourteen of the 22 genes that were differentially expressed between tolerant and resistant strains that are located in QTL regions were associated with tolerance. For the suggestive association with tolerance found on Chr 6, C-type lectin domain family 2 member H (*Clec2h*) was the most highly differentially expressed gene between tolerant and resistant strains. *Clec2h* (also known as C-type lectin-related (*Clr-f*)) is expressed in ileum, kidney, liver and in IL2-activated natural killer cells [50, 73], and plays a role in immune surveillance of the intestinal lumen [48]. We show that *Clec2h* is also expressed in spleen. *Clec2h* may facilitate bacterial detection in the spleens in tolerant mice, and thus helping them mount an appropriate immune response to STm. Alternatively, detection of *Clec2h* could result from a large influx of natural killer cells to the spleen in response to the presence of STm. Further work may identify the cell types expressing *Clec2h* during STm infection.

*Fga*, *Fgb*, and *Fgg* are three of the most highly up-regulated genes in the spleens of resistant mice and are associated with reduced spleen colonization and the QTL on Chr 3, *Scq1*. Fibrinogen is important in countering initial bacterial colonization by forming a matrix that traps bacteria and by recruiting and activating immune cells [42]. Fibrinogen kinetics (formation and degradation) are also vital to wound healing, our histopathology analysis suggests the presence of micro-abscesses in both spleen and liver that require healing [42]. Fibrinogen is also important in bacterial defense. Several bacterial species successfully modify fibrinogen to improve their proliferation, particularly *S. aureus*, supporting an evolutionary battle between fibrinogen and bacteria [74]. STm infections can alter kinetics of thrombus formation (clots) in the spleen and liver, further supporting an interaction between fibrinogen and STm [75]. In our work, the fibrinogen genes were some of the most highly differentially expressed genes, suggesting that high fibrinogen expression is a previously unknown resistance mechanism.

We had previously defined and characterized a group of Collaborative Cross strains that are highly susceptible to acute STm infection and identified several regions of the mouse genome involved [37]. In the current work, we have identified multiple phenotypes in response to long-term STm infection: delayed susceptibility, tolerance, and resistance, and characterized several hallmarks of these phenotypes. STm tolerant CC strains have lower core body temperature pre-infection, and higher, more persistent bacterial colonization than resistant strains with minimal clinical signs of disease. Disruption of the core body temperature diurnal rhythm of tolerant animals is a very early sign of disease, while gross disruptions in locomotor activity occur with a >24 hour lag. During long term infection, tolerant strains develop and do not repair damage in spleen and liver, maintain neutrophilia, have elevated levels of circulating MCP-1 and IFN-γ, and a reduction in ENA-78 relative to resistant strains. High expression of *Clec2h*, located on the QTL on Chr 6, is correlated with tolerance and the mechanistic basis underlying this association remains to be explored. Finally, we show that multiple fibrinogen genes on the QTL *Scq1* on Chr 3 are highly expressed in resistant strains, potentially revealing a new pathway involved in resistance to *Salmonella* infection.

## Materials and Methods

### Bacterial strains and media

The *Salmonella enterica* ser. Typhimurium strain (HA420) used for this study was derived from ATCC14028. HA420 is a fully virulent, spontaneous nalidixic acid resistant derivative of ATCC14028 [76]. Strains were routinely cultured in Luria-Bertani (LB) broth and plates, supplemented with antibiotics when needed at 50 mg/L nalidixic acid (Nal).

For murine infections, strains were grown aerobically at 37°C to stationary phase in LB broth with nalidixic acid and diluted to generate an inoculum of 2-5 x 10^7^ organisms in 100 microliters. Bacterial cultures used as inoculum were serially diluted and plated to enumerate colony forming units (CFU) to determine the exact titer.

### Murine strains

Collaborative cross (CC) mice were used in these experiments. All Collaborative Cross strains were obtained from UNC’s Systems Genetics Core Facility (SGCF) and either used directly for experiments or subsequently bred independently at Division of Comparative Medicine at Texas A&M University prior to these experiments Our experiments utilized 18 strains of CC mice, 3 females and 3 males per strain for a total of 112 mice (CC045 only had 2 females due to an experimental complication) (Table S5). Mice were fed Envigo Teklad Global 19% Protein Extruded Rodent Diet (irradiated, 2919) or Envigo Teklad Rodent Diet (8604) based on strain need.

### Ethics statement

All animal experiments were conducted in accordance with the Guide for the Care and Use of Laboratory Animals, and with the approval of the TAMU Institutional Animal Care and Use Committee (IACUC) under Animal use Protocol numbers: AUP# 2018-0488 D and 2015-0315 D.

### Placement of telemetry devices

5 to 9-week-old mice (CC) were anesthetized with isoflurane anesthesia. The abdomen was opened with a midline abdominal incision (up to 2 cm). Starr Life Sciences G2 E-mitter devices were loosely sutured to the ventral abdominal wall as previously described [37] to continuously monitor core body temperature and activity. Implanted mice were group-housed and monitored twice daily for signs of pain and to ensure wound closure for 7 days post-surgery. Any animals found to have serious complications after surgery were humanely euthanized. Clips were removed at 7 days post-surgery.

### Infection with *Salmonella* Typhimurium

After 7 days of acclimation in the BSL-2 facility, 8 to 12-week-old implanted CC mice were weighed and infected by gavage with a dose of 2-5 x 10^7^ CFU of *S.* Typhimurium HA420 in 100 microliters of LB broth as previously described [37]. Infected mice were monitored twice daily for signs of disease and activity by visual inspection. When telemetry data and health condition data suggested the development of clinical disease from infection, mice were humanely euthanized. If animals remained clinically healthy throughout the duration of the experiment, they were humanely euthanized at 21 days post-infection.

### Bacterial load determination

Mice were humanely euthanized by CO_2_ asphyxiation, and the spleen, liver, ileum, cecum, colon, Peyer’s patches, and mesenteric lymph nodes were collected. A third of each organ was collected in 3 mL of ice-cold PBS, weighed, homogenized, serially diluted in 1X PBS, and plated on Nal plates for enumeration of S. Typhimurium in each organ. Peyer’s patches and mesenteric lymph nodes were collected whole. Data are expressed as CFU/g of tissue.

### Telemetry monitoring

Prior to placing implanted mice on telemetry platforms, mice were moved into individual cages and provided with a cardboard hut and bedding material. Individual cages containing implanted, uninfected mice were placed onto ER4000 receiver platforms, and the collection of body temperature (once per minute), and gross motor activity (continuous measurement summed each minute) data were initiated. Body temperature and gross motor activity data were collected for 7 days from uninfected mice. Mice were removed briefly from the receiver platforms for infection and then were placed back on the platforms and data collection was resumed. Infected animals were continuously monitored by telemetry in addition to twice daily visual monitoring.

### Identification of deviation from circadian pattern of body temperature

Additional clinical information, such as the time of inoculation relative to the start of the experiment (denoted T), was used in centering time series for comparison between mice (typically, seven days after the beginning of monitoring). Quantitative detection of deviation from the baseline “off-pattern,” using temperature data were calculated on an individual basis. A temperature time series was filtered, a definition of healthy variation was defined, then the time of first “off-pattern” was calculated using that definition on post-inoculation data.

Each mouse time series was pre-processed using a moving median filter with a one day window. For a specific minute t, the median collection of temperature values from [t-720, t+720] was used in calculating a median for the value t. After this processing, healthy variation was defined as any temperature falling within the range of minimum to maximum values during the pre-inoculation phase [T-5760,T] (5760 minutes=4 days. This choice allowed for enough data to account for natural inter-day variation due to potential factors such as inter-strain variation, epigenetic differences, and sex differences, while avoiding bias due to observed acclimation time after transfer to a new facility in some mice in the first few days of observation (Fig S5A).

Identifying post-inoculation off-pattern behavior was done by identifying temperature values that fall outside the interval of healthy variation. The post-inoculation interval ranges from [T+60,T+30240] (21 days), where the one hour gap was used to avoid false positive detection due to the physical disturbance associated with inoculation (Fig S5A).

### Detection of deviation from circadian pattern of activity

While activity data do exhibit circadian patterns, this data necessitated a different approach to preprocessing compared to temperature data because activity values are inherently non-negative and have a modal value of zero. Two approaches were taken. One was from the perspective of parameter estimation of iid sampling from a statistical distribution. The second approach was based on plainly calculating the fraction of non-zero activity values measured in a moving window. Hence, we approached the analysis of activity data from the perspective of determining the parameters of a stochastic process. In the first approach, for a given time interval t, we worked from the assumption that the number of activity values observed to be i obeys the following distribution:

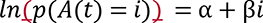

where the coefficient β, expected to be negative, corresponds to the modeling assumption that the relative drop in observed activity counts ought to decay (note 0<e^β<1 if β is negative). For instance, if the value of β= −0.693, so that e^β^ ≈1\/2, this would represent an assumption that there are half as many activity values observed to be 1 (one movement per minute) than 2 (two movements per minute). In actuality, this decay coefficient is typically seen to be β≈-0.025 to β≈-0.015, representing that observed activity values decay by half around every 27-46 values.

This theory is implemented in practice by windowing a mouse’s activity time series, creating a binning (empirical distribution) (*i*,*c*(*i*)),, the performing a log-linear fit by transforming *c*(*i*)→*ln*(1 + *c*(*i*)) and applying linear least-squares regression to calculate α and β. Statistical models with simpler assumptions stemming from a “memoryless” assumption were attempted but did not see any feasible agreement with observed activity data (Fig S5B).

The second approach mentioned above produced a qualitatively cleaner signal, which again is based on windowing a mouse’s activity time series, and calculating the fraction of activity values in a window for which A(t)>0. One might expect not weighting by intensity of activity loses information, but we saw this approach to filtering activity data resulted in a time series for which we could apply the same methods as with the temperature time series (Fig S5B)

### Histopathology

After euthanasia, samples of liver, spleen, ileum, cecum, and colon from each mouse were collected and fixed in 10% neutral buffered formalin at room temperature for 24 hours, stored in 70% ethanol before embedding in paraffin, sectioning at 5 µm, and staining with hematoxylin and eosin (H&E). Histologic sections of all tissues were evaluated in a blinded manner by bright field microscopy and scored on a scale from 0 to 4 for tissue damage by a board-certified veterinary pathologist (Table S4). The combined scores of spleens and liver from individual mice were used to calculate the median and interquartile range for each group. Whole slide images of liver and spleen H&E-stained sections were captured as digital files by scanning at 40x using a 3DHistech Pannoramic SCAN II FL™ scanner (Epredia, Kalamazoo, MI). Digital files were processed by Aiforia Hub (Aiforia, Cambridge, MA) software for generating the images with 100 µm scale bars.

### Complete blood count

Whole blood was collected by cheek bleed for complete blood count (CBC) one week prior to surgery for the pre-infection sample. Blood from infected animals was collected at necropsy by cardiac puncture. Blood was collected from additional uninfected control animals by cardiac puncture and median values were used as the baseline for mice that did not serve as their own pre-infection control. All blood was collected into EDTA tubes and analyzed on an Abaxis VetScan HM5.

### Cytokine and chemokine analysis

Serum was stored at −80°C and thawed on ice immediately prior to cytokine assays. Serum cytokine levels were evaluated using an Invitrogen ProcaratPlex Cytokine/Chemokine Convenience Panel 1A 36 plex kit as per manufacturer instructions (ThermoFisher). Briefly, magnetic beads were added to each well of the 96-well plate and washed on the Bio-Plex Pro Wash Station. 25uL of samples and standards were then added to the plate along with 25uL of universal assay buffer. Plates were shaken at room temperature for 1 hour and washed. 25uL of detection antibody was added an incubated for 30 minutes and washed away, followed by 50uL of SAPE for 30 minutes and washed away. 120uL of reading buffer was added and plates were evaluated using a Bio-Plex 200 (BioRad). Samples and standards were assayed in duplicate, and samples were diluted as needed to get 25ul of serum per duplicate. The kit screens for the following 36 cytokines and chemokines: IFN gamma; IL-12p70; IL-13; IL-1 beta; IL-2; IL-4; IL-5; IL-6; TNF alpha; GM-CSF; IL-18; IL-10; IL-17A; IL-22; IL-23; IL-27; IL-9; GRO alpha; IP-10; MCP-1; MCP-3; MIP-1 alpha; MIP-1 beta; MIP-2; RANTES; Eotaxin; IFN alpha; IL-15/IL-15R; IL-28; IL-31; IL-1 alpha; IL-3; G-CSF; LIF; ENA-78/CXCL5; M-CSF. Median values of uninfected animals for each strain were used as the baseline for that strain.

### QTL analysis

gQTL, an online resource designed specifically to identify CC QTLs, was used to find putative QTL associations [77]. Briefly, median values of each strain for various parameters were uploaded to the website and QTL analysis was performed using 1000 permutations with “automatic” transformation. Automatic transformation picks either log or square root transformations, whichever best normalizes the data.

### Transcriptomic analysis

Spleen tissue was snap frozen in liquid nitrogen and stored in −80°C freezer until ribonucleic acid (RNA) sequencing could be performed. All molecular work was performed in the Molecular Genomics Core of the Texas A&M Institute for Genome Sciences and Society (TIGSS). The following protocol was adapted from the Molecular Genomics Core, and TIGSS personnel aided in data acquisition and plot generation. Spleens were homogenized in Trizol. RNA samples were quantified with a Qubit Fluorometer (Life Technologies) with a broad range RNA assay and concentrations were normalized for library preparation. RNA quality was verified on an Agilent TapeStation with a broad range RNA ScreenTape. Total RNA sequencing libraries were prepared using the Illumina TruSeq Stranded mRNA-seq preparation kit. Barcoded libraries were pooled at equimolar concentrations and sequenced on an Illumina NovaSeq 6000 2×150 S4 flow cell. RNA-seq libraries were trimmed to remove adapter sequences and low-quality bases using TrimGalore version 0.6.6, with Cutadapt version 3.0 and FastQC version 0.11.9.

The Colorado State University team performed data processing of the RNAseq using a pipeline consisting of STAR alignment [78], FeatureCounts [79], then DeSeq2 R package was used for differential gene expression analysis [80]. The STAR alignment and FeatureCounts were done using mouse genome version GRCm38 ensembl version 100 available from Ensembl [81]. All infected samples were normalized to their uninfected control samples (or an average uninfected control if no strain specific samples were available). These strain values were then grouped by infection response and a tolerant or resistant mean was used for comparisons between the groups. Tolerant was then set as the baseline, so genes with positive values were up-regulated in resistant strains and down-regulated in tolerant strains, while genes with negative values were down-regulated in resistant strains and up-regulated in tolerant strains.

---

**Supplementary Figure 1: Ileum, cecum, and colon CFU/g are not significantly different between tolerant and resistant strains.** A. Ileum, B. Cecum, and C. Colon CFU per gram of organ at the time of necropsy. Each circle represents an individual mouse and medians and interquartile ranges are indicated. T = tolerant (open circles), R = resistant (closed circles). 11 strains included in the graph.

**Supplementary Figure 2: Pre-infection activity levels are not significantly different between groups.** Fraction of time active (0-1) for resting, 24-hour, and active periods for delayed susceptible (DS, filled triangles), tolerant (T, open circles), and resistant (R, filled circles) mice. Median and interquartile range indicated.

**Supplementary Figure 3: Ileum, cecum, and colon are minimally damaged**. Ileum and cecum/colon histopathology mean and standard deviation shown for each group.

**Supplementary Figure 4: Susceptible strains have the most damage at one week post-infection compared to the other groups.** The mean of spleen and liver histopathology scores for tissue damage and standard deviation indicated. T = tolerant (black bars), R = resistant (gray bars), DS = delayed susceptible (red bars), S = susceptible (yellow bar) 1W = one week post-infection (filled bars). Kruskal-Wallis test was performed to find significant differences (* *P* <0.05, ** *P* <0.01, *** *P* <0.001, **** *P* <0.0001).

**Supplemental Figure 5: Examples of telemetry calculations**. Illustrations of how A. temperature and B. activity calculations were determined. Specifically maximum, median, minimum pre-infection temperature and activity as well “off-pattern” for post-infection.

**Supplemental Figure 6: Liver and spleen colonization is significantly lower in resistant strains three weeks post-infection.** A. Liver and B. spleen CFU per gram of organ at the time of necropsy. Each circle represents an individual mouse and medians and interquartile ranges are indicated. T = tolerant (open circles), R = resistant (closed circles), 1W = one week post-infection, 3W = three weeks post-infection. 11 strains included in the graph. Kruskal-Wallis test was performed to find significant differences (* *P* <0.05, ** *P* <0.01, *** *P* <0.001, **** *P* <0.0001).

**Supplemental Table 1: Median values of core body temperature for rest period, 24-hour period, and active period.**

**Supplemental Table 2: Complete Blood Counts from infected mice taken at necropsy minus uninfected mean for respective strain for 18 CC strains.** Mean and standard deviation shown. See Excel document.

**Supplemental Table 3: 36 Cytokines from infected mice taken at necropsy minus uninfected mean for respective strain for 18 CC strains**. Mean and standard deviation shown. See Excel document.

**Supplemental Table 4: Histopathology scoring matrix with descriptions and pictures.** Final score per organ corresponds to whichever section has the highest score. A. Scoring matrix from Dr. L. Garry Adams. B. Representative images of spleen and liver histology scoring matrix. Tissues were sectioned and stained with H&E before being analyzed. Images from Dr. L. Garry Adams.

**Supplementary Table 5: Number of mice used and facility origin for 18 CC strains.**

## References

1. Roy MF, Malo D. Genetic regulation of host responses to Salmonella infection in mice. Genes Immun. 2002;3: 381–393. doi:10.1038/sj.gene.6363924

2. Santos RL, Tükel Ç, Raffatellu M, Bäumler AJ, Tsolis RM, Adams LG, et al. Life in the inflamed intestine, Salmonella style. Trends Microbiol. 2009;17: 498–506. doi:10.1016/j.tim.2009.08.008

3. Majowicz SE, Musto J, Scallan E, Angulo FJ, Kirk M, O’Brien SJ, et al. The global burden of nontyphoidal salmonella gastroenteritis. Clin Infect Dis. 2010;50: 882– 889. doi:10.1086/650733

4. Gilchrist JJ, MacLennan C a., Hill AVS. Genetic susceptibility to invasive Salmonella disease. Nat Rev Immunol. 2015;15: 452–463. doi:10.1038/nri3858

5. Stanaway JD, Parisi A, Sarkar K, Blacker BF, Reiner RC, Hay SI, et al. The global burden of non-typhoidal salmonella invasive disease: a systematic analysis for the Global Burden of Disease Study 2017. Lancet Infect Dis. 2019;19: 1312–1324. doi:10.1016/S1473-3099(19)30418-9

6. Gordon MA. Salmonella infections in immunocompromised adults. J Infect. 2008;56: 413–422. doi:10.1016/j.jinf.2008.03.012

7. Keestra-gounder AM, Tsolis RM, Bäumler AJ. Now you see me, now you don’t : the interaction of Salmonella with innate immune receptors. Nat Rev Microbiol. 2015;13: 206–216. doi:10.1038/nrmicro3428

8. Lalmanach AC, Montagne A, Menanteau P, Lantier F. Effect of the mouse Nramp1 genotype on the expression of IFN-γ gene in early response to Salmonella infection. Microbes Infect. 2001;3: 639–644. doi:10.1016/S1286-4579(01)01419-8

9. Arpaia N, Godec J, Lau L, Sivick KE, McLaughlin LM, Jones MB, et al. TLR signaling is required for salmonella typhimurium virulence. Cell. 2011;144: 675– 688. doi:10.1016/j.cell.2011.01.031

10. Mastroeni P, Ugrinovic S, Chandra A, MacLennan C, Doffinger R, Kumararatne D. Resistance and susceptibility to Salmonella infections: Lessons from mice and patients with immunodeficiencies. Rev Med Microbiol. 2003;14: 53–62. doi:10.1097/00013542-200304000-00002

11. Caron J, Loredo-Osti JC, Laroche L, Skamene E, Morgan K, Malo D. Identification of genetic loci controlling bacterial clearance in experimental Salmonella enteritidis infection: An unexpected role of Nramp1 (Slc11a1) in the persistence of infection in mice. Genes Immun. 2002;3: 196–204. doi:10.1038/sj.gene.6363850

12. Sancho-Shimizu V, Malo D. Sequencing, Expression, and Functional Analyses Support the Candidacy of Ncf2 in Susceptibility to Salmonella Typhimurium Infection in Wild-Derived Mice . J Immunol. 2006;176: 6954–6961. doi:10.4049/jimmunol.176.11.6954

13. Khan RT, Chevenon M, Yuki KE, Malo D. Genetic dissection of the Ity3 locus identifies a role for Ncf2 co-expression modules and suggests Selp as a candidate gene underlying the Ity3.2 locus. Front Immunol. 2014;5: 1–13. doi:10.3389/fimmu.2014.00375

14. Li Q, Cherayil BJ. Role of toll-like receptor 4 in macrophage activation and tolerance during Salmonella enterica serovar Typhimurium infection. Infect Immun. 2003;71: 4873–4882. doi:10.1128/IAI.71.9.4873-4882.2003

15. Bihl F, Larivière L, Qureshi ST, Flaherty L, Malo D. LPS-hyporesponsiveness of mnd mice is associated with a mutation in Toll-like receptor 4. Genes Immun. 2001;2: 56–59. doi:10.1038/sj.gene.6363732

16. Churchill G, Airey D, Allayee H, … The Collaborative Cross, a community resource for the genetic analysis of complex traits. Nat Rev Genet. 2004;36: 1133–1137.

17. Aylor DL, Valdar W, Foulds-mathes W, Buus RJ, Verdugo R a, Baric RS, et al. Genetic analysis of complex traits in the emerging Collaborative Genetic analysis of complex traits in the emerging Collaborative Cross. Genome Res. 2011;21: 1213–1222. doi:10.1101/gr.111310.110

18. Threadgill DW, Miller DR, Churchill G a, de Villena FP-M. The collaborative cross: a recombinant inbred mouse population for the systems genetic era. ILAR J. 2011;52: 24–31. doi:10.1093/ilar.52.1.24

19. Iraqi FA, Mahajne M, Salaymah Y, Sandovski H, Tayem H, Vered K, et al. The genome architecture of the collaborative cross mouse genetic reference population. Genetics. 2012;190: 389–401. doi:10.1534/genetics.111.132639

20. Philippi J, Xie Y, Miller DR, Bell TA, Zhang Z, Lenarcic AB, et al. Using the emerging Collaborative Cross to probe the immune system. Genes Immun. 2014;15: 38–46. doi:10.1038/gene.2013.59.Using

21. Abu Toamih Atamni H, Nashef A, Iraqi FA. The Collaborative Cross mouse model for dissecting genetic susceptibility to infectious diseases. Mamm Genome. 2018;29: 471–487. doi:10.1007/s00335-018-9768-1

22. Smith CM, Sassetti CM. Modeling Diversity: Do Homogeneous Laboratory Strains Limit Discovery? Trends Microbiol. 2018;26: 892–895. doi:10.1016/j.tim.2018.08.002

23. Korth MJ, Tchitchek N, Benecke A, Katze MG. Systems approach to influenza-virus host interactions and the pathogenesis of highly virulent and pandemic viruses. Semin Immunol. 2013;25. doi:10.1007/s11103-011-9767-z.Plastid

24. Rasmussen AL, Okumura A, Ferris MT, Green R, Feldmann F, Kelly SM, et al. Host genetic diversity enables Ebola hemorrhagic fever pathogenesis and resistance. Science (80-). 2014;346: 987–991. doi:10.1126/science.1259595.Host

25. Price A, Okumura A, Haddock E, Feldmann F, Meade-White K, Sharma P, et al. Transcriptional Correlates of Tolerance and Lethality in Mice Predict Ebola Virus Disease Patient Outcomes. Cell Rep. 2020;30: 1702–1713.

26. Plant JE, Glynn AA. Genetic control of resistance to Salmonella typhimurium infection in high and low antibody responder mice. Clin Exp Immunol. 1982;50: 283–290.

27. Kover PX, Schaal BA. Genetic variation for disease resistance and tolerance among Arabidopsis thaliana accessions. Proc Natl Acad Sci. 2002;99: 11270– 11274. doi:10.1073/pnas.102288999

28. Howick VM, Lazzaro BP. Genotype and diet shape resistance and tolerance across distinct phases of bacterial infection. BMC Evol Biol. 2014;14: 1–13. doi:10.1186/1471-2148-14-56

29. Merkling SH, Bronkhorst AW, Kramer JM, Overheul GJ, Schenck A, Van Rij RP. The Epigenetic Regulator G9a Mediates Tolerance to RNA Virus Infection in Drosophila. PLoS Pathog. 2015;11: 1–25. doi:10.1371/journal.ppat.1004692

30. Louie A, Song KH, Hotson A, Thomas Tate A, Schneider DS. How Many Parameters Does It Take to Describe Disease Tolerance? PLoS Biol. 2016;14: 1– 21. doi:10.1371/journal.pbio.1002435

31. Raberg L, Sim D, Read AF. Disentangling Genetic Variation for Resistance and Tolerance to Infectious Disease in Animals. Science (80-). 2007;318: 812–814. doi:10.1126/science.1229223

32. Read AF, Graham AL, Råberg L. Animal defenses against infectious agents: Is damage control more important than pathogen control? PLoS Biol. 2008;6: 2638– 2641. doi:10.1371/journal.pbio.1000004

33. Medzhitov R, Schneider DS, Soares MP, Caldwell RM, Schafer JF, Compton LE, et al. Disease tolerance as a defense strategy. Science. 2012;335: 936–41. doi:10.1126/science.1214935

34. Martins R, Carlos AR, Braza F, Thompson JA, Bastos-Amador P, Ramos S, et al. Disease Tolerance as an Inherent Component of Immunity. Annu Rev Immunol. 2019;37: 405–437. doi:10.1146/annurev-immunol-042718-041739

35. Ayres JS, Schneider DS. Two ways to Survive an Infection: What Resistance and Tolerance Can Teach Us About Treatments for Infectious Diseases. Nat Rev Immunol. 2008;8: 889–895. doi:10.1038/nri2432.Two

36. Malo D, Skamene E. Genetic control of host resistance to infection. Trends Genet. 1994;10: 365–371. doi:10.1016/0168-9525(94)90133-3

37. Scoggin K, Lynch R, Gupta J, Nagarajan A, Sheffield M, Elsaadi A, et al. Genetic background influences survival of infections with Salmonella enterica serovar Typhimurium in the Collaborative Cross. PLoS Genet. 2022;accepted.

38. Deshmane SL, Kremlev S, Amini S, Sawaya BE. Monocyte chemoattractant protein-1 (MCP-1): An overview. J Interf Cytokine Res. 2009;29: 313–325. doi:10.1089/jir.2008.0027

39. Ingram JP, Brodsky IE, Balachandran S. Interferon–γ in Salmonella Pathogenesis: New Tricks for an Old Dog. Cytokine. 2017;98: 27–32. doi:10.1016/j.cyto.2016.10.009.Interferon

40. Koltsova EK, Ley K. The mysterious ways of the chemokine cxcl5. Immunity. 2010;33: 7–9. doi:10.1016/j.immuni.2010.07.012

41. Mei J, Liu Y, Dai N, Favara M, Greene T, Jeyaseelan S, et al. CXCL5 regulates chemokine scavenging and pulmonary host defense to bacterial infection. Immunity. 2010;33: 106–117. doi:10.1016/j.immuni.2010.07.009

42. Kearney KJ, Ariëns RAS, Macrae FL. The Role of Fibrin(ogen) in Wound Healing and Infection Control. Semin Thromb Hemost. 2021. doi:10.1055/s-0041-1732467

43. Ravasi T, Hsu K, Goyette J, Schroder K, Yang Z, Rahimi F, et al. Probing the S100 protein family through genomic and functional analysis. Genomics. 2004;84: 10–22. doi:10.1016/j.ygeno.2004.02.002

44. O’Keefe EJ, Hamilton EH, Lee SC, Steinertt P. Trichohyalin: A structural protein of hair, tongue, nail, and epidermis. J Invest Dermatol. 1993;101. doi:10.1016/0022-202X(93)90503-A

45. Abdelhamed Z, Lukacs M, Cindric S, Ali S, Omran H, Stottmann RW. A novel hypomorphic allele of Spag17 causes primary ciliary dyskinesia phenotypes in mice. DMM Dis Model Mech. 2020;13. doi:10.1242/DMM.048645

46. Andrade AD, Almeida PGC, Mariani NAP, Freitas GA, Kushima H, Filadelpho AL, et al. Lipopolysaccharide-induced epididymitis modifies the transcriptional profile of Wfdc genes in mice. Biol Reprod. 2021;104: 144–158. doi:10.1093/biolre/ioaa189

47. Rutkowski E, Leibelt S, Born C, Friede ME, Bauer S, Weil S, et al. Clr-a: A Novel Immune-Related C-Type Lectin-like Molecule Exclusively Expressed by Mouse Gut Epithelium. J Immunol. 2017;198: 916–926. doi:10.4049/jimmunol.1600666

48. Leibelt S, Friede ME, Rohe C, Gütle D, Rutkowski E, Weigert A, et al. Dedicated immunosensing of the mouse intestinal epithelium facilitated by a pair of genetically coupled lectin-like receptors. Mucosal Immunol. 2015;8: 232–242. doi:10.1038/mi.2014.60

49. Pahari S, Negi S, Aqdas M, Arnett E, Schlesinger LS, Agrewala JN. Induction of autophagy through CLEC4E in combination with TLR4: an innovative strategy to restrict the survival of Mycobacterium tuberculosis. Autophagy. 2020;16: 1021– 1043. doi:10.1080/15548627.2019.1658436

50. Zhang Q, Rahim MMA, Allan DSJ, Tu MM, Belanger S, Abou-Samra E, et al. Mouse Nkrp1-Clr Gene Cluster Sequence and Expression Analyses Reveal Conservation of Tissue-Specific MHC-Independent Immunosurveillance. PLoS One. 2012;7. doi:10.1371/journal.pone.0050561

51. Finch S, Shoemark A, Dicker AJ, Keir HR, Smith A, Ong S, et al. Pregnancy zone protein is associated with airway infection, neutrophil extracellular trap formation, and disease severity in bronchiectasis. Am J Respir Crit Care Med. 2019;200: 992–1001. doi:10.1164/rccm.201812-2351OC

52. Krause K, Azouz F, Nakano E, Nerurkar VR, Kumar M. Deletion of pregnancy zone protein and murinoglobulin-1 restricts the pathogenesis of West Nile virus infection in mice. Front Microbiol. 2019;10: 1–9. doi:10.3389/fmicb.2019.00259

53. Van De Steeg E, Wagenaar E, Van Der Kruijssen CMM, Burggraaff JEC, De Waart DR, Oude Elferink RPJ, et al. Organic anion transporting polypeptide 1a/1b-knockout mice provide insights into hepatic handling of bilirubin, bile acids, and drugs. J Clin Invest. 2010;120: 2942–2952. doi:10.1172/JCI42168

54. Hagenbuch B, Meier PJ. Organic anion transporting polypeptides of the OATP/SLC21 family: Phylogenetic classification as OATP/SLCO super-family, new nomenclature and molecular/functional properties. Pflugers Arch Eur J Physiol. 2004;447: 653–665. doi:10.1007/s00424-003-1168-y

55. Monack DM, Bouley DM, Falkow S. Salmonella typhimurium Persists within Macrophages in the Mesenteric Lymph Nodes of Chronically Infected Nramp1+/+ Mice and Can Be Reactivated by IFNγ Neutralization. J Exp Med. 2004;199: 231– 241. doi:10.1084/jem.20031319

56. Prior KF, O’Donnell AJ, Rund SSC, Savill NJ, van der Veen DR, Reece SE. Host circadian rhythms are disrupted during malaria infection in parasite genotype-specific manners. Sci Rep. 2019;9: 1–12. doi:10.1038/s41598-019-47191-8

57. Huitron-Resendiz S, Marcondes MCG, Flynn CT, Lanigan CMS, Fox HS. Effects of simian immunodeficiency virus on the circadian rhythms of body temperature and gross locomotor activity. Proc Natl Acad Sci U S A. 2007;104: 15138–15143. doi:10.1073/pnas.0707171104

58. Lough G, Kyriazakis I, Bergmann S, Lengeling A, Doeschl-Wilson AB. Health trajectories reveal the dynamic contributions of host genetic resistance and tolerance to infection outcome. Proc R Soc B Biol Sci. 2015;282. doi:10.1098/rspb.2015.2151

59. Okada K, Yano M, Doki Y, Azama T, Iwanaga H, Miki H, et al. Injection of LPS Causes Transient Suppression of Biological Clock Genes in Rats. J Surg Res. 2008;145: 5–12. doi:10.1016/j.jss.2007.01.010

60. Gschwandtner M, Derler R, Midwood KS. More Than Just Attractive: How CCL2 Influences Myeloid Cell Behavior Beyond Chemotaxis. Front Immunol. 2019;10: 1–29. doi:10.3389/fimmu.2019.02759

61. Bossink AWJ, Paemen L, Jansen PM, Hack CE, Thijs LG, Van Damme J. Plasma levels of the chemokines monocyte chemotactic proteins-1 and -2 are elevated in human sepsis. Blood. 1995;86: 3841–3847. doi:10.1182/blood.v86.10.3841.bloodjournal86103841

62. Van Coillie E, Van Damme J, Opdenakker G. The MCP/eotaxin subfamily of CC chemokines. Cytokine Growth Factor Rev. 1999;10: 61–86. doi:10.1016/S1359-6101(99)00005-2

63. Martin GS. Sepsis, severe sepsis and septic shock: changes in incidence, pathogens and outcomes. Expert Rev Anti Infect Ther. 2012;10: 701–706. doi:10.1016/S0140-6736(18)30696-2

64. Chaudhry H, Zhou J, Zhong Y, Ali MM, McGuire F, Nagarkatti PS, et al. Role of Cytokines as a Double-edged Sword in Sepsis. PMC. 2015;27: 669–684.

65. Rutledge BJ, Rayburn H, Rosenberg R, North RJ, Gladue RP, Corless CL, et al. High level monocyte chemoattractant protein-1 expression in transgenic mice increases their susceptibility to intracellular pathogens. J Immunol. 1995;155: 4838–43. Available: http://www.ncbi.nlm.nih.gov/pubmed/7594486

66. Feng WX, Flores-Villanueva PO, Mokrousov I, Wu XR, Xiao J, Jiao WW, et al. CCL2-2518 (A/G) polymorphisms and tuberculosis susceptibility: A meta-analysis. Int J Tuberc Lung Dis. 2012;16: 150–156. doi:10.5588/ijtld.11.0205

67. Hui WW, Hercik K, Belsare S, Alugubelly N, Clapp B, Rinaldi C, et al. Salmonella enterica serovar Typhimurium alters the extracellular proteome of macrophages and leads to the production of proinflammatory exosomes. Infect Immun. 2018;86. doi:10.1128/IAI.00386-17

68. Eckmann L, Kagnoff MF. Cytokines in host defense against Salmonella. Microbes Infect. 2001;3: 1191–1200. doi:10.1016/S1286-4579(01)01479-4

69. Rowell DL, Eckmann L, Dwinell MB, Carpenter SP, Raucy JL, Yang SK, et al. Human hepatocytes express an array of proinflammatory cytokines after agonist stimulation or bacterial invasion. Am J Physiol - Gastrointest Liver Physiol. 1997;273.

70. Raberg L, Graham AL, Read AF. Decomposing health: tolerance and resistance to parasites in animals. Philos Trans R Soc B Biol Sci. 2009;364: 37–49. doi:10.1098/rstb.2008.0184

71. McCarville JL, Ayres JS. Disease tolerance: concept and mechanisms. Curr Opin Immunol. 2018;50: 88–93. doi:10.1016/j.coi.2017.12.003

72. Ram R, Mehta M, Balmer L, Gatti DM, Morahan G. Rapid identification of major-effect genes using the collaborative cross. Genetics. 2014;198: 75–86. doi:10.1534/genetics.114.163014

73. Plougastel B, Dubbelde C, Yokoyama WM. Cloning of Clr, a new family of lectin-like genes localized between mouse Nkrp1a and Cd69. Immunogenetics. 2001;53: 209–214. doi:10.1007/s002510100319

74. Ko YP, Flick MJ. Fibrinogen Is at the Interface of Host Defense and Pathogen Virulence in Staphylococcus aureus Infection. Semin Thromb Hemost. 2016;42: 408–421. doi:10.1055/s-0036-1579635

75. Beristain-Covarrubias N, Perez-Toledo M, Flores-Langarica A, Zuidscherwoude M, Hitchcock JR, Channell WM, et al. Salmonella-induced thrombi in mice develop asynchronously in the spleen and liver and are not effective bacterial traps. Blood. 2019;133: 600–604. doi:10.1182/blood-2018-08-867267

76. Bogomolnaya LM, Santiviago CA, Yang HJ, Baumler AJ, Andrews-Polymenis HL. “Form variation” of the O12 antigen is critical for persistence of Salmonella Typhimurium in the murine intestine. Mol Microbiol. 2008;70: 1105–1119. doi:10.1111/j.1365-2958.2008.06461.x

77. Konganti K, Ehrlich A, Rusyn I, Threadgill D. gQTL : A Web Application for QTL Analysis Using the Collaborative Cross Mouse Genetic Reference Population. Genes Genomes Genet. 2018;8. doi:10.1534/g3.118.200230

78. Dobin A, Davis CA, Schlesinger F, Drenkow J, Zaleski C, Jha S, et al. STAR: Ultrafast universal RNA-seq aligner. Bioinformatics. 2013;29: 15–21. doi:10.1093/bioinformatics/bts635

79. Liao Y, Smyth GK, Shi W. FeatureCounts: An efficient general purpose program for assigning sequence reads to genomic features. Bioinformatics. 2014;30: 923– 930. doi:10.1093/bioinformatics/btt656

80. Love MI, Huber W, Anders S. Moderated estimation of fold change and dispersion for RNA-seq data with DESeq2. Genome Biol. 2014;15: 1–21. doi:10.1186/s13059-014-0550-8

81. Howe KL, Achuthan P, Allen J, Allen J, Alvarez-Jarreta J, Ridwan Amode M, et al. Ensembl 2021. Nucleic Acids Res. 2021;49: D884–D891. doi:10.1093/nar/gkaa942

